# Profiling cellular heterogeneity and fructose transporter expression in the rat nephron by integrating single-cell and microdissected tubule segment transcriptomes

**DOI:** 10.1101/2023.12.20.572656

**Authors:** Ronghao Zhang, Darshan Aatmaram Jadhav, Benjamin Kramer, Agustin Gonzalez-Vicente, the Kidney Precision Medicine Project

**Author notes:** Corresponding author: Agustin Gonzalez-Vicente 1: Department of Physiology and Biophysics Case Western Reserve University, 10900 Euclid Avenue, Robbins-E532 Cleveland, OH-44106. Darshan Aatmaram Jadhav and Ronghao Zhang contributed equally to this manuscript.

## Abstract

Single-cell RNA sequencing (scRNAseq) is a crucial tool in kidney research. These technologies cluster cells according to transcriptome similarity, irrespective of the anatomical location and ordering within the nephron. Thus, a cluster transcriptome may obscure heterogeneity of the cell population within a nephron segment. Elevated dietary fructose leads to salt-sensitive hypertension, in part by fructose reabsorption in the proximal tubule (PT). However, organization of the four known fructose transporters in apical PTs (SGLT4, SGLT5, GLUT5 and NaGLT1) remains poorly understood. We hypothesized that cells within each subsegment of the proximal tubule exhibit complex, heterogenous fructose transporter expression patterns. To test this hypothesis we analyzed rat and kidney transcriptomes and proteomes from publicly available scRNAseq and tubule microdissection databases. We found that microdissected PT-S1 segments consist of 81±12% cells with scRNAseq-derived transcriptional characteristics of S1, whereas PT-S2 express a mixture of 18±9% S1, 58±8% S2, and 19±5% S3 transcripts, and PT-S3 consists of 75±9% S3 transcripts. The expression of all four fructose transporters was detectable in all three PT segments, but key fructose transporters SGLT5 and GLUT5 progressively increased from S1 to S3, and both were significantly upregulated in S3 vs. S1/S2 (Slc5a10: 1.9 log_2_FC, p<1×10^-299^; Scl2a5: 1.4 log_2_FC, p<4×10^-105^). A similar distribution was found in human kidneys. These data suggest that S3 is the primary site of fructose reabsorption in both humans and rats. Finally, because of the multiple scRNAseq transcriptional phenotypes found in each segment our findings also imply that anatomic labels applied to scRNAseq clusters may be misleading.

## Introduction

Single-cell sequencing (scRNAseq) technologies cluster cells according to transcriptome similarity, without consideration for the anatomical arrangement, which is critical for kidney tubular epithelial cell functions. Such clusters are associated with distinct segments of the nephron based on the expression of marker genes characteristic of those segments. However, given the potential for cellular diversity within each segment, a transcriptional cluster comprised only of cells with highly similar transcriptional phenotypes may not provide an accurate representation of the entire cell population within a specific region of the nephron. Thus, to explore the association between single-cell cluster transcriptional phenotypes and anatomical localization, we conducted two analyses, which compare transcriptomes from manually microdissected rat nephron segments to transcriptomes derived from dispersed cells and scRNAseq methods available in the public domain.

Modern Western diets are rich in sodium chloride (NaCl) and fructose or fructose-containing syrups (1), while in Asian countries such as China consumption of sugar-sweetened beverages has doubled within the last decade (2, 3). As such, the average calories ingested from fructose in industrialized countries, expressed as a percentage of the recommended average caloric intake exceeds 10% (4–6). Importantly, individuals consuming more than 74 g/day of fructose, equivalent to ∼15% of calories on a 2000 kcal/day diet, have higher blood pressure (7). Previous studies have shown that rodents consuming 10 to 20% of their calories as free-fructose develop salt-sensitive hypertension within a week (8–12). In addition, in chronic models of metabolic syndrome, such as those where experimental diets contain nearly 50% of dietary fructose by weight, restricting Na^+^ intake ameliorates the increase in blood pressure (11, 13). These findings provide a connection between the ingestion of fructose and salt and the development of hypertension.

Fructose undergoes glomerular filtration, and the proximal tubule (PT) reabsorbs the bulk of it. Four apical transporters with vastly different kinetic properties could transport fructose: SGLT5 (fructose k_m_: 0.62 mmol/l), NAGLT1 (km: 7.8 mmol/l), SGLT4 (k_m_>10 mmol/l) and GLUT5 (km: 12.6 mmol/l). Some of these transporters also exhibit the ability to transport glucose, for which the two monosaccharide may compete (**Table S1**). Furthermore, the distribution of fructose transporters varies along the three subsegments of the PT (14), for which the contribution of each subsegment to overall fructose transport remains poorly understood. The PT is also the only nephron segment expressing fructokinase, an intracellular enzyme that phosphorylates fructose, thereby sequestering it within the cell (15, 16). Thus, unlike glucose, which is not metabolized by PTs and returned into the blood, fructose is broken down into 3-carbon intermediates, which are then metabolized by glycolysis or gluconeogenesis, or synthesized to diacylglycerol and neutral lipids (17). Importantly, fructokinase consumes ATP and is not regulated by substrate or product availability (15, 16). Excessive intracellular fructose could therefore deplete ATP, and lead to inflammation and endothelial injury (18, 19).

Our study challenges the dogma that fructose is uniformly reabsorbed according to nephron segment. Rather, we hypothesize that cells within each subsegment of the proximal tubule exhibit complex, heterogeneous fructose transporter expression patterns, which impact the fructose transport capacity. To explore this hypothesis, we extensively analyzed publicly available resources and integrated kidney single-cell transcriptomes with bulk transcriptomic data from microdissected kidney regions.

## Methods

### Data analysis

Unless otherwise noted, all data were analyzed in R: A language and environment for statistical computing and graphics (https://www.R-project.org/). Some data inspection, cleaning or formatting were conducted in Notepad++ (https://notepad-plus-plus.org/) or in Microsoft Excel.

### Rat kidney single-cell transcriptomes

Rat whole kidney single-cell RNA sequencing transcriptomes (scRNAseq) were obtained from the Gene Expression Omnibus (GSE137869). We used filtered_features_bc_matrix files from three pools of male (GSM4331828, GSM4331829, GSM4331830) and three pools of female (GSM4331831, GSM4331832, GSM4331833) Sprague-Dawley rats. To generate these matrices, sequences from the microfluidic droplet platform were de-multiplexed and aligned to the rat genome (Rnor_6.0) using CellRanger (2.2.1) with default parameters (20). Standard protocols were implemented in Seurat (4.3.0) for quality control and data analysis. In brief, a Seurat object was created from each filtered_features_bc_matrix after selecting features detected in at least three cells and cells containing at least 200 features. All six objects were merged yielding 19414 cells with 16821 features. Then the following filters were applied: nFeature_RNA > 560, nFeature_RNA < 4500, nCount_RNA < 30000, percent.MT < 40, percent.Ribosomal < 30, percent.Largest.Gene < 25. Filtered data were log-normalized. Cell-cycle scores were calculated using the cc.genes.updated.2019 genes and regressed during data scaling (21, 22). DoubletFinder (2.0.3) was used to remove doublets with parameters optimized with the function *paramSweep_v3()*. The entire quality control process removed ∼30% of observations, yielding 16821 features across 13596 cells. After quality control, individual runs were split and SCTransform was applied. Then 3000 integration features were identified and all datasets were integrated using the *IntegrateData()* function. Principal Component Analysis (PCA) identified 30 relevant dimensions for downstream processing. The function *FindClusters()* identified 36 clusters at a resolution of 1.6. The function *FindMarkers()* with the Wilcox test was used to identify cluster markers. Pseudobulk transcriptomes were obtained using the function *AverageExpression()*.

### Human kidney single-nucleus transcriptomes

An h5Seurat file containing single-nucleus RNA sequencing (snRNAseq) data was downloaded from the Kidney Precision Medicine Project (KPMP) tissue atlas (atlas.kpmp.org) on 04/13/2023 using the following filters: Experimental Strategy: Single-cell RNA-Seq, Workflow Type: Aggregated Clustered Data, File Name: c798e11b-bbde-45dd-bd91-487f27c93f8f_WashU-UCSD_HuBMAP_KPMP-Biopsy_10X-R_12032021.h5Seurat. The file contains aggregated cluster data including 30395 features across 110346 cells as described by Lake et al. (23). In the KPMP Seurat object the aggregated clusters are annotated by: ***1) “class”*** (cell Class) i.e. Epithelial, Stromal, Immune, Endothelial or Neural, ***2) “subclass.l1”*** (subRegion) i.e. podocytes (POD), parietal epithelial cells (PEC), PT, descending thin limb (DTL), ascending thin limb (ATL), thick ascending limb (TAL), distal convoluted tubule (DCT) connecting tubule (CNT), principal cell (PC) and intercalated cell (IC), Immune (IMN), fibroblasts (FIB), neural (NEU), papillary (Pape) and vascular smooth muscle/pericyte (VSM/P), and Endothelial (EC) cells, and ***3) “subclass.l2”*** (cell type). Pseudobulk transcriptomes were obtained using the function AverageExpression() for cell Class and subRegion.

### Rat microdissected tubule segments transcriptomes

To map single-cell transcriptional phenotypes with spatial and structural features of the nephron, publicly available bulk transcriptomes (PRJNA24440) from rat microdissected nephron segments (34) were downloaded from the National Center for Biotechnology Information (NCBI) Sequence Read Archive using SRA Toolkit 3.0.2. All raw sequencing FASTQ files were reprocessed using command lines in Ubuntu 22.04. In brief, quality control was performed on all files using FastQC v0.11.9 (24), and low-quality reads were trimmed using Trimmomatic v0.39 (25). The remaining sequences were aligned to the rat genome obtained from the NCBI RefSeq Database (GCF_015227675.2_mRatBN7.2_genomic.fna.gz) using Burrows-Wheeler Aligner (BWA) v0.7.17-r1188 (26). The alignment was then sorted and exported by Samtools v1.13 (27) in the format of BAM files. Reads were counted from BAM files using the R package Rsubread v2.12.0 with a reference to genome annotation obtained from RefSeq (GCF_015227675.2_mRatBN7.2_genomic.gtf). The raw-counts matrices and metadata indicating the anatomical location of microdissected segments were processed and incorporated into a DESeq2 object (DESeq2 v1.34.0) (28). Sequencing depth was normalized using the median of ratio method.

### Correlation analysis

Pseudobulk transcriptomes from individual rat scRNAseq clusters, regions, and cell types were contrasted with different transcriptomes from rats and humans, as well as with rat tubular proteomics data. For these comparisons we used genomewide Pearson correlation, as implemented by the function *cor(x, method = “Pearson”, use = “pairwise.complete.obs”)* in R. For visualizations, we used the function *pheatmap(x)* from the R package pheatmap_1.0.12.

### Transcriptional clusters identity assignment

The genomewide Pearson correlation between pseudobulk transcriptomes from rat clusters and KPMP snRNAseq cell classes was used to identify and extract epithelial cells (**Figure 1**). Clusters were manually assigned to different kidney cell types using marker genes from the HuBMAP Kidney v1.2 cell markers (29) dataset, and other sources (30) (**Figure 2**). Clusters 17 and 27 presented low and diffuse expression of tubular epithelial cells markers (**Figure S1**) and were therefore excluded from subsequent analysis. No cluster expressed markers of glomerular visceral epithelial cell (podocyte) markers was low in all clusters, while the expression of PEC markers was scattered across different clusters (**Figure S2**), for which we assumed a very low count of glomerular epithelial cells.

**Figure 1:**
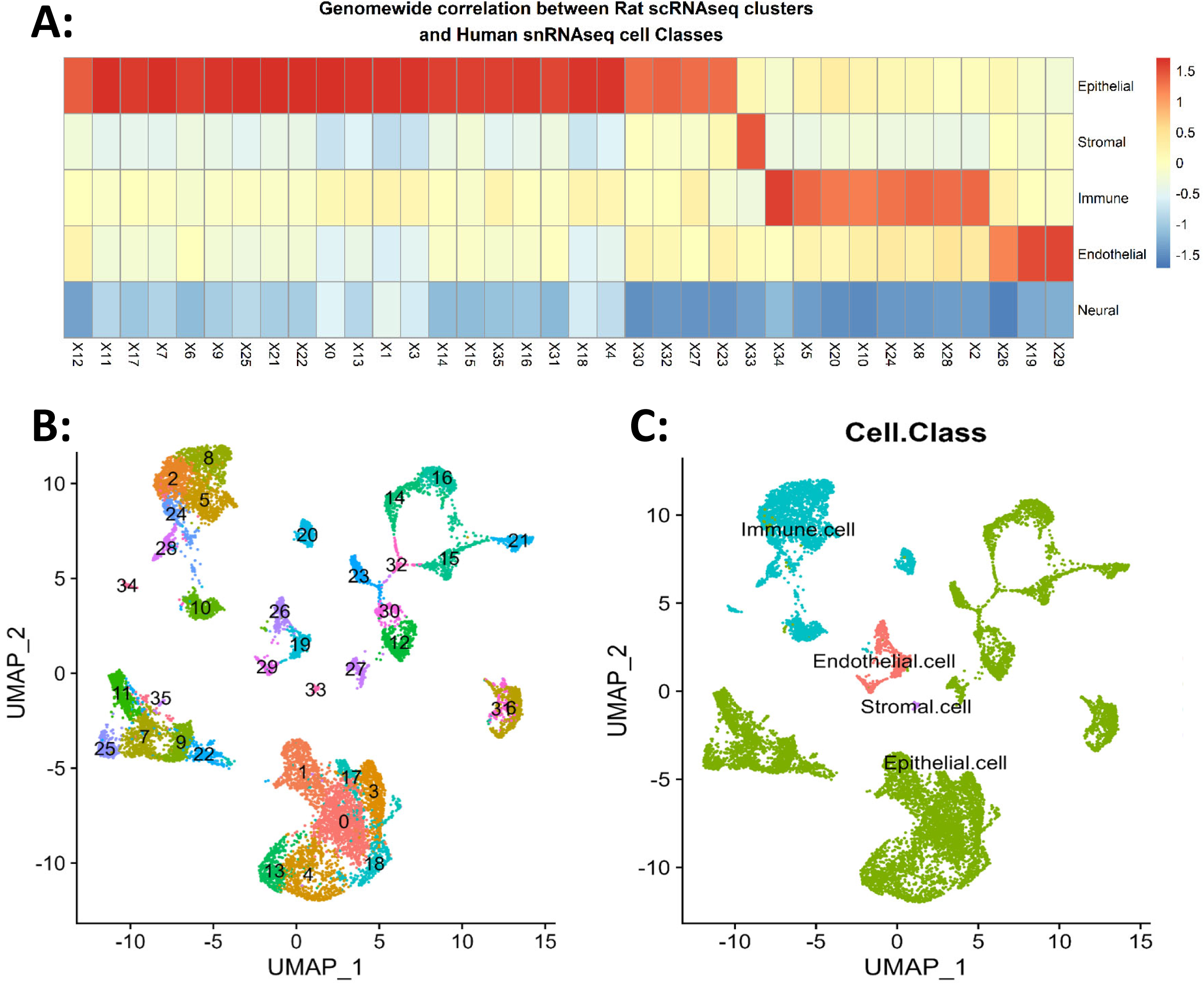
All rat scRNAseq clusters identified in the rat kidney were assigned to different cell classes by conducting a genomewide Pearson correlation with pseudobulk transcriptomes from human snRNAseq cell classes. **A)** Normalized Pearson correlation coefficient between rat scRNAseq clusters and humans snRNAseq cell classes yielded: 1) Epithelial, 24 clusters; 2) Stromal, 1 cluster; 3) Immune, 8 clusters; and 4) Endothelial, 3 clusters, while no cluster in the rat dataset correlated with neural cells. **B)** UMAP projection of individual clusters. **C)** UMAP projection of the four assigned cell classes.

**Figure 2:**
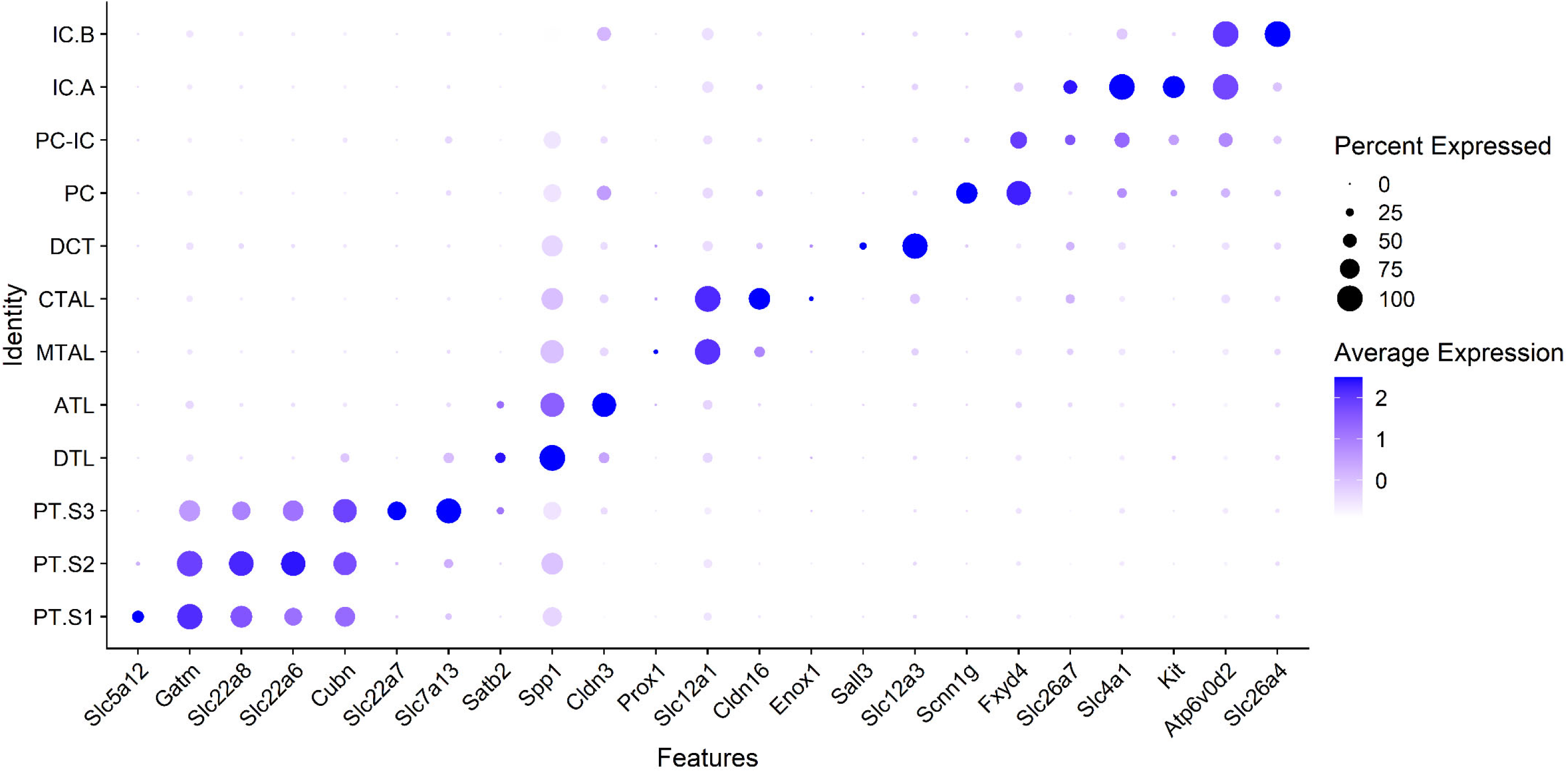
Cluster assignments using HuBMAP Kidney v1.2 cell markers.

### Rat microdissected tubule segments transcriptomes deconvolution

We utilized CibersortX (31), to deconvolute bulk transcriptomes from microdissected nephron segments using the scRNAseq clusters generated here as a reference. This approach estimated the fractional composition of distinct transcriptional phenotypes within the bulk tubule segment transcriptomes. Raw sequencing files from both datasets were downloaded from the Sequence Read Archive (SRA) and reprocessed as described above. The number of detected genes was 34,322 for the bulk RNAseq data and 15,282 for the scRNAseq pseudobulk transcriptomes, with a shared gene space of 13,373 that was used for deconvolution. CibersortX was run with predetermined parameters, and without permutations for significance testing to focus solely on estimating cell type fractions.

### Rat microdissected tubule segments proteomics

Quantitative proteomics from rat PT subsegments (32) were downloaded from the National Heart, Lung, and Blood Institute (NHLBI-NIH) Epithelial Systems Biology Laboratory (33).

## Results

### Rat kidney scRNAseq map

Using publicly available scRNAseq transcriptomes from rat whole kidneys, we identified 36 cell clusters, which were assigned to different cell classes using the KPMP snRNAseq reference data as follows: Epithelial, 24 clusters; Stromal, 1 cluster; Immune, 8 clusters; and Endothelial, 3 clusters, while no cluster in the rat dataset correlated with neural cells (**Figures 1A and 1C**). Then, the tubular epithelial cell cluster identities were annotated using a curated list of cell markers from HuBMAP Kidney v1.2 (29) and other sources (23, 30) (**Figure S1**). We identified and annotated clusters of most tubular epithelial cell types including PT.S1, PT.S2, PT.S3, DTL, ATL, medullary TAL (MTAL), cortical TAL (CTAL), DCT, PC, intercalated cells types A and B (IC.A and IC.B, respectively) (**Figures 1B and 2**). Cluster 32 (**Figures 1B**), corresponded to an undefined cell type expressing markers from both, principal and intercalated cells (PC.IC). In addition, we contrasted cluster regions to the snRNAseq subregional transcriptomes obtained from the KPMP, demonstrating similarity with human tubular cell annotations (**Figure 3.B**). Of note, the rat scRNAseq dataset contains nearly 10% of the cell count of the KPMP snRNAseq and with less sequencing depth. As such the 12 tubular epithelial clusters identified in the rat, provided enough resolution for comparison with the subRegion (subclass.l1) but not the cell type (subclass.l2) identified in humans.

**Figure 3:**
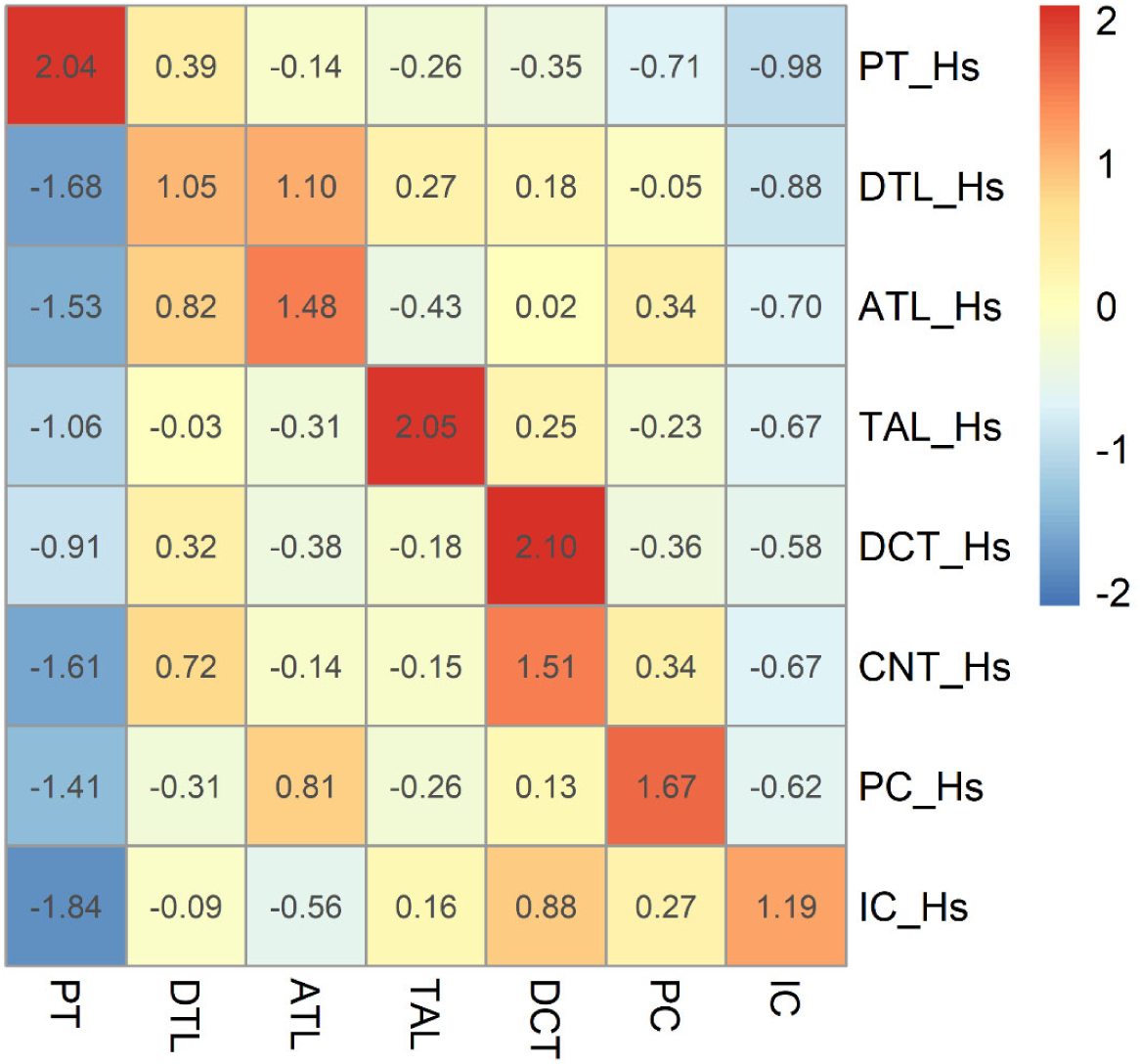
Correlation between rat scRNAseq cell cluster regions and transcriptomes from human kidneys. Normalized Pearson correlation between rat cell clusters and human single-nucleus (sn)RNAseq sub-regional transcriptomes.

PCA of all cell types found in the dataset, including epithelial, endothelial, stromal and immune, shows that the tubular epithelium separates from other cell classes along the first principal component (PC1) (**Figure 4A**), while PT cells separate from all other epithelial cells along the second principal component (PC2) (**Figure 4B**). Moreover, the distribution of tubular epithelial cells along PC2 resembles the anatomy of the nephron with the more distal cells exhibiting the greater separation from PT cells. To focus our analysis on the tubular epithelium, we extracted epithelial cells for downstream analyses.

**Figure 4:**
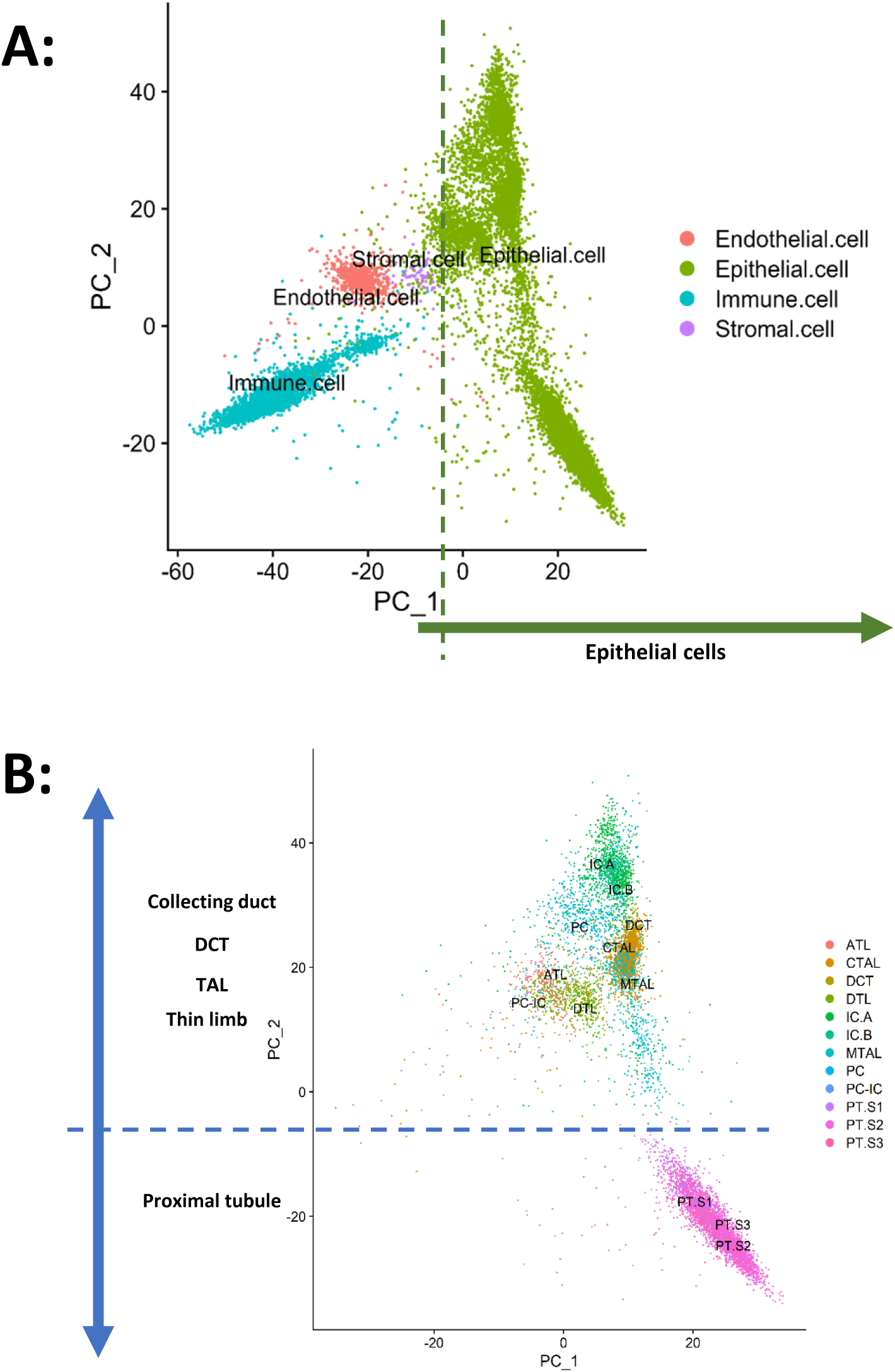
Principal component analysis of rat kidney cells. **A)** Epithelial cells separate from other cell classes along PC1, and **B)** while tubular epithelial cells separate along PC2 resembling the anatomy of the nephron.

Firstly, we measured the correlation between cluster average transcriptomes and bulk transcriptomes in manually microdissected tubules. We used the “Median” gene expression of each segment as originally published by Knepper and collaborators (34). The use of the median gene expression value is robust against contamination with other segments as far as such contamination is present in less than 50% of the samples. We found four well-defined nephron regions presenting the highest correlation with their corresponding cell clusters: 1) PTs, 2) thin limbs, 3) TAL/DCT and 4) CNT/CD (**Figure 5**). Bulk transcriptomics of microdissected S1 segments primarily correlate with the PT.S1 scRNAseq cluster, but also present a strong association with the PT.S2 scRNAseq cluster. Microdissected S2 segments present similar correlations with all three PT subsegment clusters. Microdissected S3 segments present the strongest correlation overall with the PT.S3 cluster. Microdissected segments of the loop of Henle present a strong correlation with the corresponding cell types characterized in the scRNAseq. Notably, microdissected DCT correlated with the DCT cluster expressing the sodium-chloride symporter (NCC, *Slc12a3*) but also with the MTAL cluster. The PC cluster dominates the correlations in CNT and collecting duct segments. Finally, the “PC.IC” cell type is most highly correlated with the inner medullary collecting duct (IMCD) segment transcriptome.

**Figure 5:**
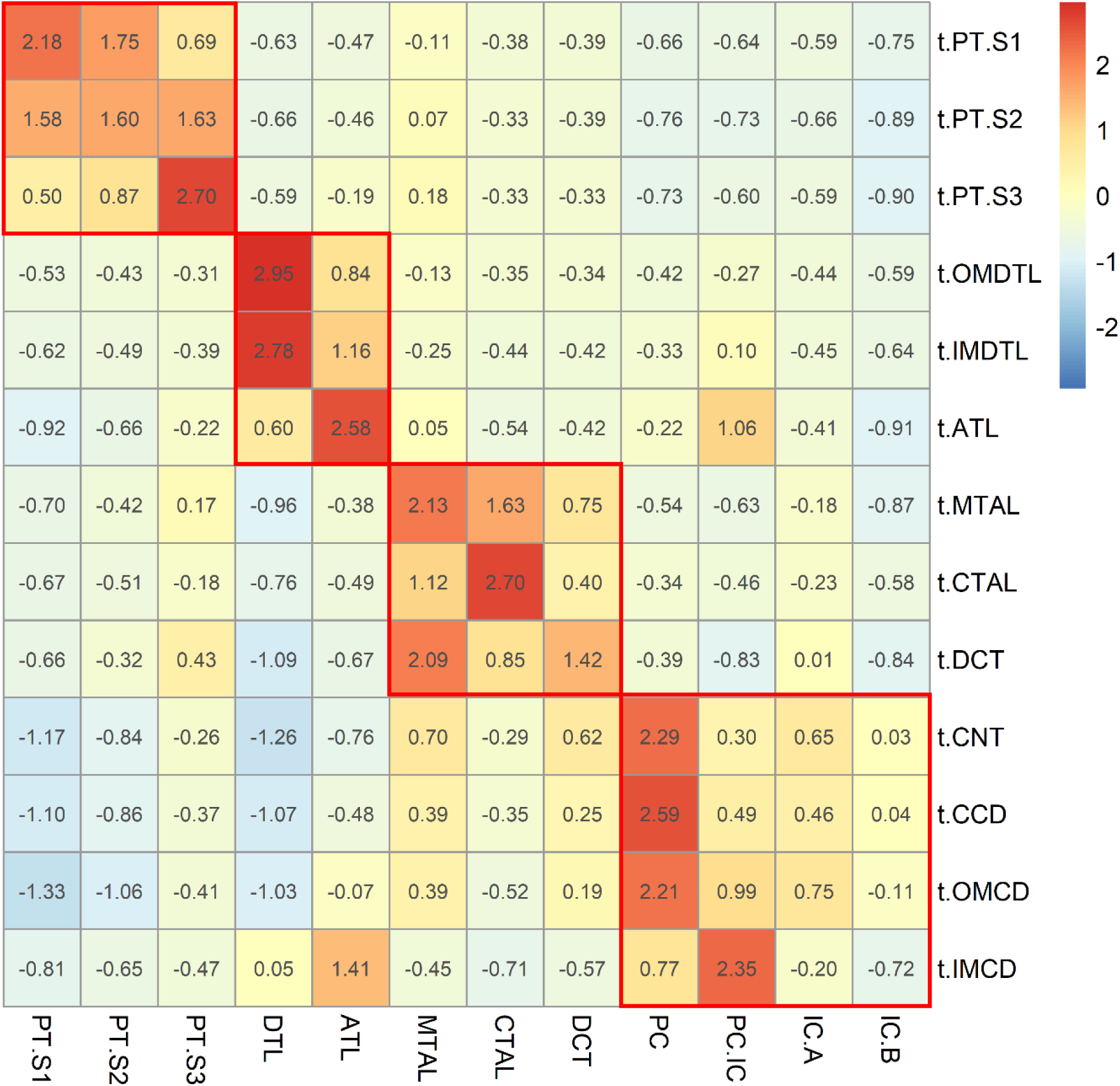
Correlation between rat scRNAseq clusters transcriptomes and bulk transcriptomes from rat microdissected nephron segments. A genomewide Pearson correlation was calculated and normalized. For each microdissected segment the values on the table represent the number of standard deviations that the correlation with each cluster deviates from the average of all clusters.

The second analysis estimated the proportion of distinct cell transcriptional phenotypes identified by scRNAseq within each segment. We used scRNAseq cluster transcriptomes as a reference to deconvolute the transcriptomes from microdissected segments. First, we reprocessed raw sequencing files to obtain the count matrix from each replicate. Unlike the first analysis, where segment-specific transcriptomes were reconstructed through median gene expression, here we calculated the mean gene expression for each segment. For this, we initially deconvoluted each individual replicate against the rat scRNAseq regional transcriptomes to identify possible contaminations. Microdissected tubule transcriptomes with either: 1) less than 88% of cells from the corresponding region, or 2) more than 10% of an adjacent single region, were dropped from downstream analysis (**Table S2_Quality Control**). The remaining 90 transcriptomes were deconvoluted against scRNAseq cellular clusters (**Table S2_Results**). Cell percentage results are presented as mean +/− standard error. We found that PT.S1 segments primarily consist of S1 cells (81 ± 12 %) with a small proportion of S2 cells (12 ± 11 %). Microdissected PT.S2 segments contain a mix of S1 (18 ± 9 %), S2 (58 ± 8 %), and S3 (19 ± 5 %) cells. PT.S3 segments were primarily composed of S3 cells (75 ± 9 %), with lower proportions of the S2 (10 ± 5 %) and S1 (8 ± 7 %) cell types. The thin portions of the Loop of Henle displayed a high percentage of the corresponding cell type: outer medullary DTL (OMDTL) (91 ± 2 %, DTL), inner medullary DTL (IMDTL) (86 ± 3 % DTL), and ATL (78 ± 9 % ATL). Microdissected MTAL segments were a mix of MTAL (25 ± 15 %) and CTAL (69 ± 15%) cells, while microdissected CTAL segments were almost exclusively composed of CTAL cells (95 ± 2 %). DCT segments exhibited a majority (80 ± 5 %) of DCT cells with some principal (2 ± 1 %) and intercalated (1 ± 1 %) cells. In CNT, CCD, and OMCD, we observed a continuous increase in the abundance of PC cells (58 ± 6 %, 75 ± 3 %, and 78 ± 6 %, respectively) along with a decrease in IC (32 ± 2 %, 20 ± 3 %, and 15 ± 4 %, respectively). Microdissected IMCD contained PC (51 ± 7 %), IC (4 ± 1%), and a significant percentage (27 ± 11 %) of a distinct cell type not found in other segments, that we previously labeled “PC-IC” (**Figure 2**). IMCD also presents a significant percentage of ATL (17 ± 4 %) cells, which may reflect contamination (**Table S2_Summary**).

### Metabolism of fructose

We next studied the expression of sugar transporters and fructose metabolic enzymes. Fructose can be produced endogenously from glucose via the polyol pathway (35, 36). We found that the expression of sorbitol dehydrogenase (*Sord*), the initial enzyme in the polyol pathway was stronger in PTs, while mRNA encoding aldose reductase (*Akr1b1*), the rate-limiting step in the polyol pathway was predominantly expressed in medullary segments (**Figure 6.A-B**). The distributions of these enzymes in rat match that in the human kidney (**Figure 6.C**) (23, 37).

**Figure 6:**
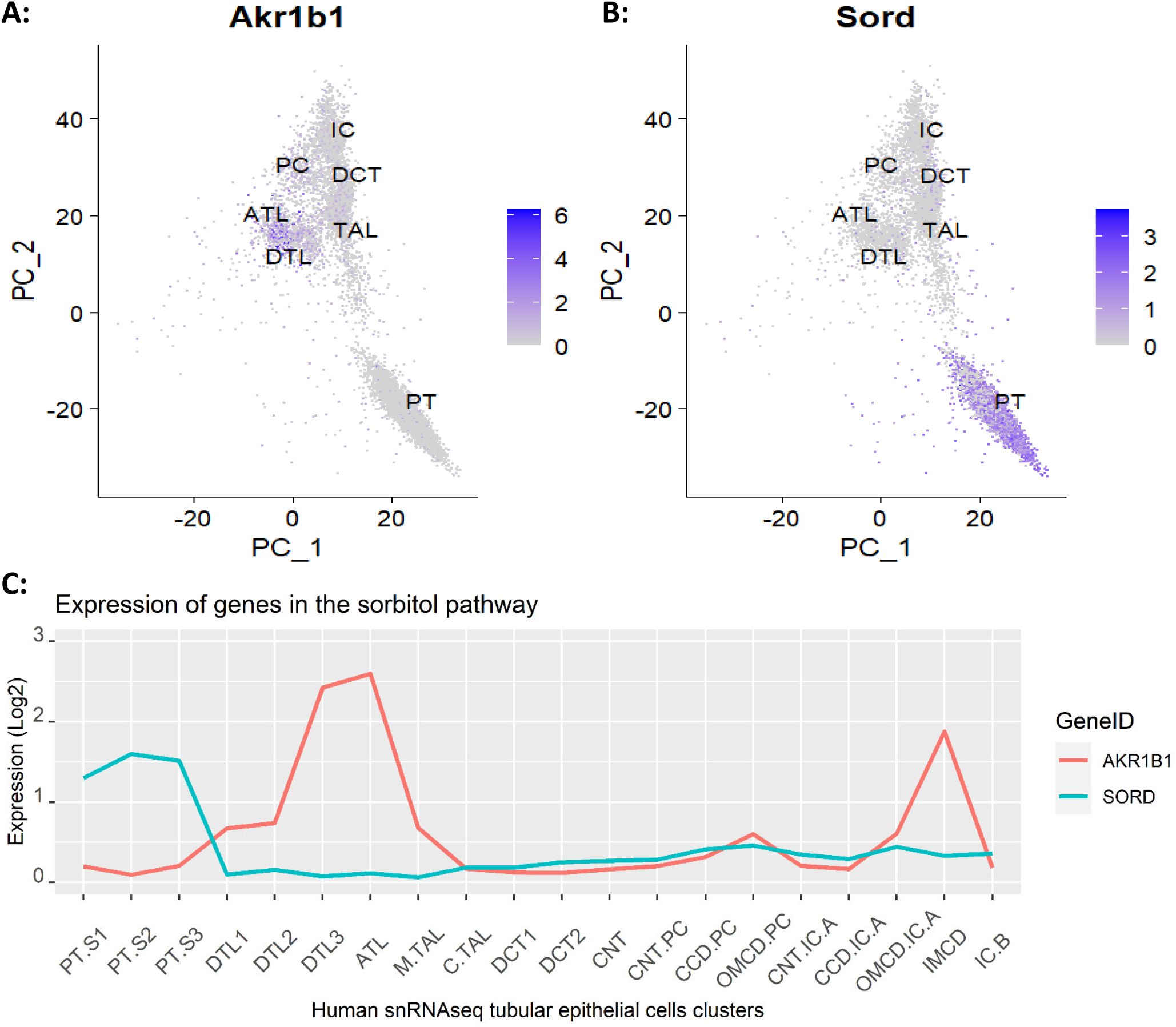
Expression of enzymes from the sorbitol pathway. **A)** Aldolase reductase (*Akr1b1*) converts glucose to sorbitol and is predominantly expressed in medullary segments. **B)** Sorbitol dehydrogenase (*Sord*) converts sorbitol to fructose, and is predominantly expressed in proximal tubules. **C)** Expression of the aldolase reductase (*AKR1B1*) and sorbitol dehydrogenase (*SORD*) genes in human kidney.

We next examined the expression of enzymes that allow fructose metabolites to enter glycolytic pathways and neutral lipids synthesis (35, 36). Fructokinase (*Khk*), which phosphorylates fructose to fructose-1-P, and triokinase (*Tkfc*) are specific enzymes in this process. **Figure 7A** shows that *Khk* and *Tkfc* are restricted to PTs, indicating that this segment metabolizes the bulk of fructose. On the contrary, aldolase B (*Aldob*) and triosephosphate isomerase (*Tpi1*) are shared with glycolysis, and the anatomic distribution is correspondingly not exclusive to PT (**Figure 7B**).

**Figure 7:**
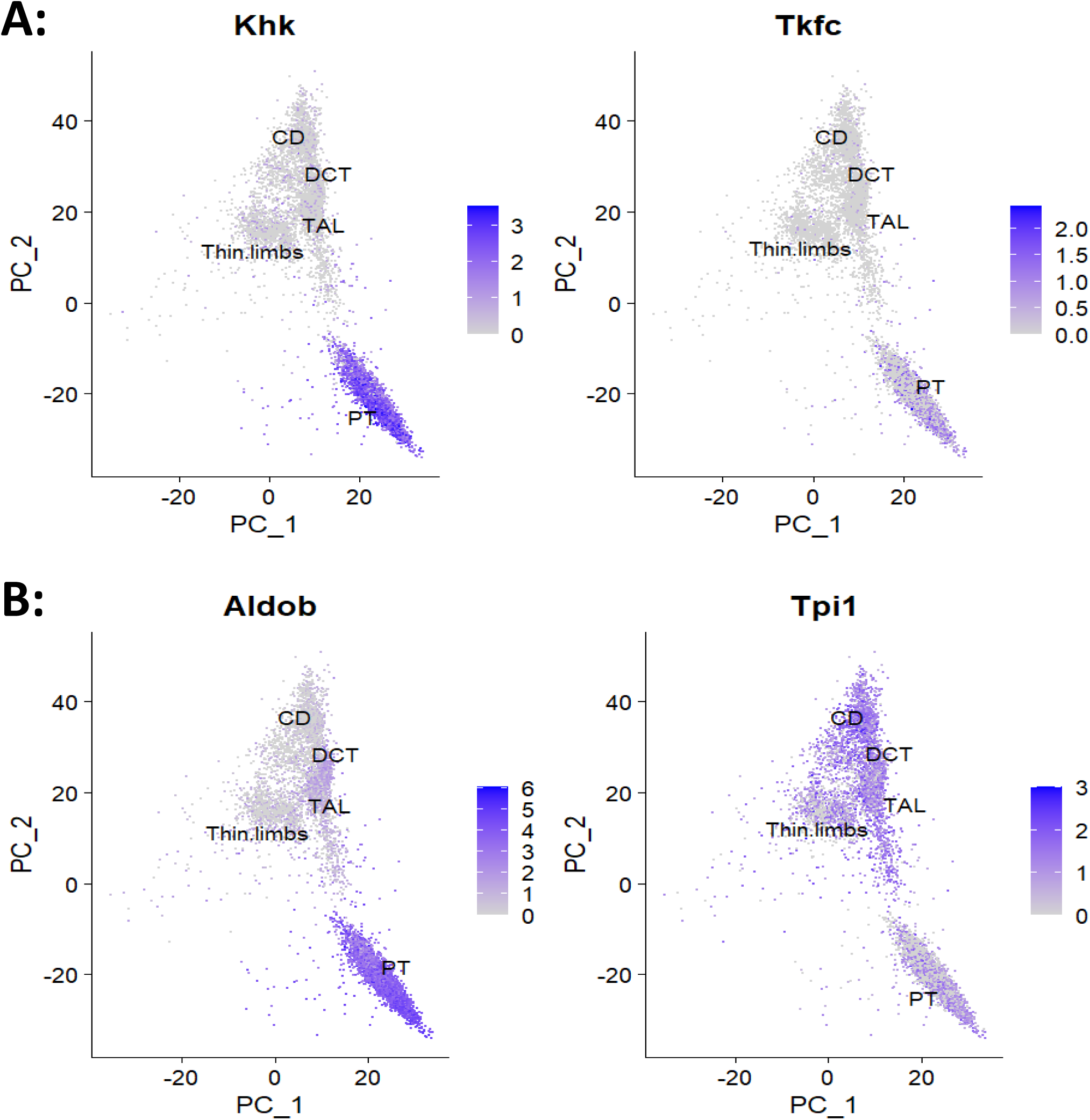
Expression of enzymes necessary to metabolize fructose into synthetic pathways. **A)** Enzymes specific to fructose metabolism, fructokinase and (*Khk*) and triokinase (*Tkfc*) are restricted to proximal tubules. **B)** Enzymes shared with glycolysis, aldolase B (*Aldob*) and triosephosphate isomerase (*Tpi1*) are widely expressed in other segments in addition to proximal tubules.

Next, we extracted the PT clusters and conducted differential gene expression analysis between PT.S1, PT.S2, and PS.S3 to identify significant differences in the expression of fructose transporters and metabolic enzymes (**Table S3**). Several genes were differentially expressed in each cluster as compared to the other two together, including fructose and glucose transporters **(Figure 8)**. Both SGLT2 (Slc5a2) and GLUT2 (Slc2a2) were upregulated in PT.S1 cluster while SGLT5 (*Slc5a10*) was downregulated. The PT.S2 cluster only presented downregulation of the SGLT2 gene (Slc5a2). Finally, SGLT5 (*Slc5a10*) and GLUT5 (*Slc2a5*) were both upregulated in PT.S3, while SGLT2 (*Slc5a2*) and NAGLT1 (*Naglt1*) were downregulated in this segment. These data indicate that the pairs SGLT5/GLUT5 and SGLT2/GLUT2 are transcriptionally paired in rat PT cells. SGLT1 (*Slc5a1*), GLUT1 (*Slc2a1*) and SGLT4 (*Slc5a9*) were not differentially expressed in any of the three PT clusters; however, there were smaller differences in expression below the significance threshold (**Figures S3**) across clusters. Genes coding for enzymes participating in the polyol pathways were not differentially expressed across clusters.

**Figure 8:**
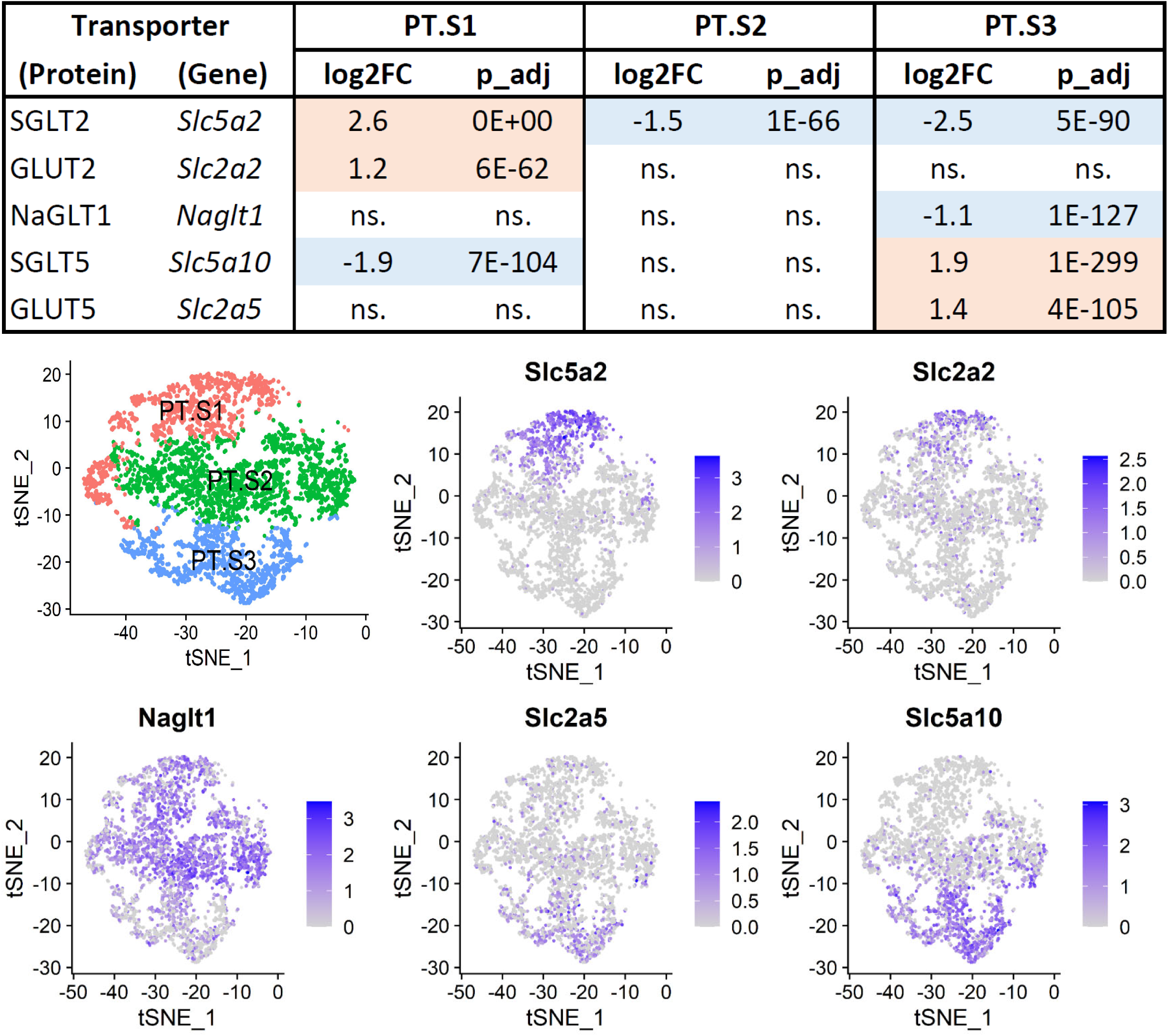
Sugar transporters differentially expressed across proximal tubule cell types. **Upper table)** log2 fold change (Log2FC) and p values on each segment, as compared to the other two segments together. **Lower panels)** TSNE projections of proximal tubule clusters showing a transcript density map of the differentially expressed sugar transporter in rat proximal tubule scRNAseq clusters. Only differentially expressed sugar transporters are shown in the figure.

Finally, to qualitatively assess whether RNA transcript abundance resembles differences in protein expression, we examined the protein expression of fructose transporters and metabolic enzymes in microdissected rat PT segments and compare them with the RNA expression in rat scRNAseq (**Figure 9**). Of note, fructose transporter mRNA and protein expression tightly paralleled one another (**Figures 9A and 9B**). Importantly, all four fructose transporters were expressed in both S2 and S3. A similar analysis of glucose transporters as well as the transcript levels in human snRNAseq data can be found in the supplement (**Figures S4 and S5**). Although subtle differences were observed between rat and human fructose transporter expression (**Figures 9A and S4A**), fructokinase and triokinase expression were remarkably similar between rat and human proximal tubule segments (**Figures 9C and S4C**).

**Figure 9:**
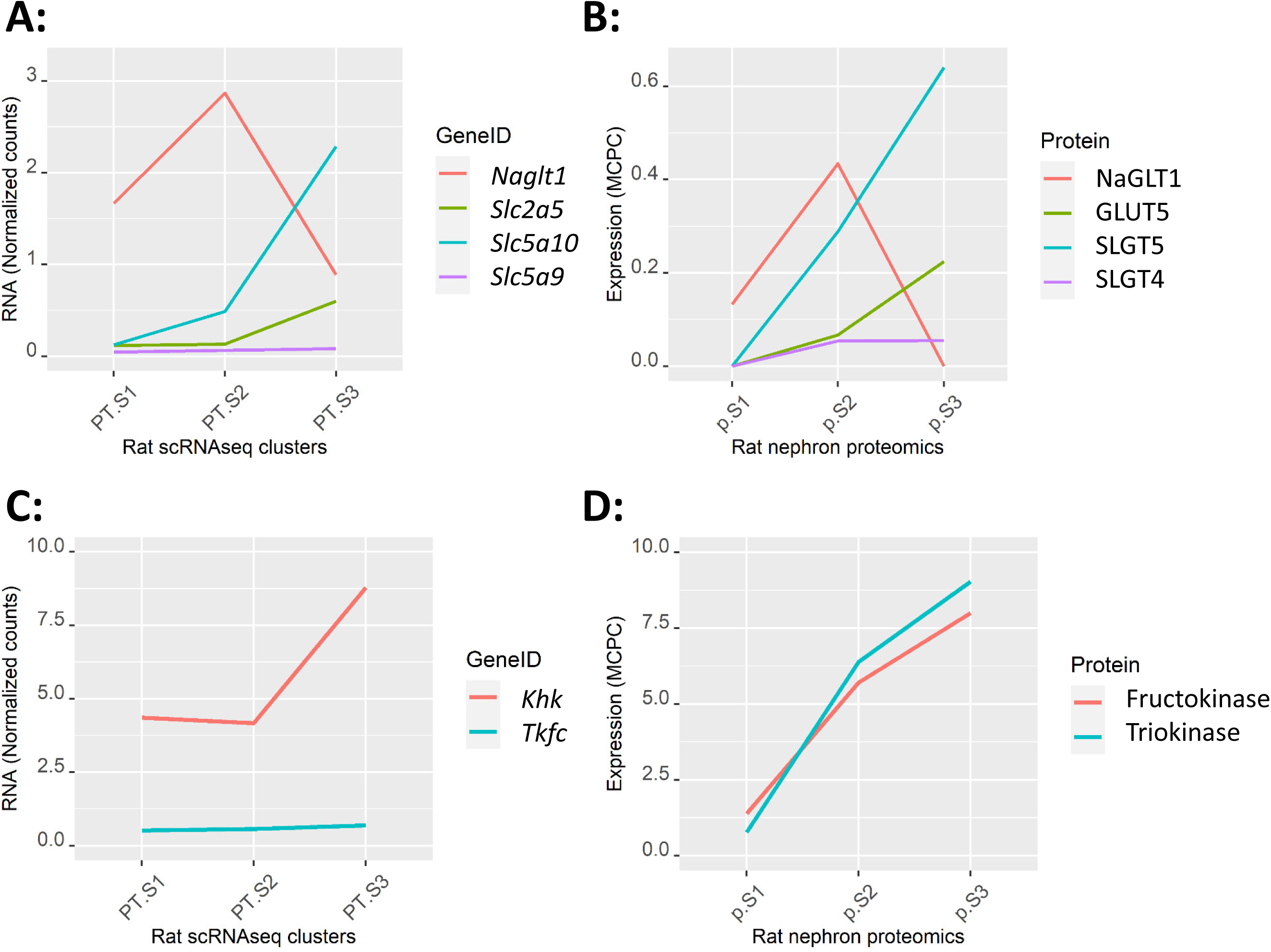
Normalized RNA transcripts of fructose transporters in rat scRNAseq proximal tubule clusters (**A**) compared to their corresponding proteins expressed in “million copies per cell” (MCPC) in microdissected rat proximal tubule segments (**B**). Normalized RNA transcripts of fructokinase (*Khk*) and Triokinase (*Tkfc*) in rat scRNAseq proximal tubule clusters (**C**) compared to their corresponding proteins expression in microdissected rat proximal tubule segments (**D**). The order and colors of GeneIDs on the left panels (**A and C**) match that of the corresponding protein on the right panels (**B and D**).

## Discussion

To study the distribution of fructose transporters and metabolic enzymes transcripts along the rat nephron, we constructed a scRNAseq map of rat kidney cells. To identify different cell types, we used human kidney transcriptomes from the KPMP and HuBMAP kidney cell markers as a reference (**Figures 1**-**3**). Thus, our annotations of rat kidney cells align with those found in human kidneys. The map encompasses immune, stromal, and endothelial cells, as well as most tubular epithelial cell types including PT.S1, PT.S2, PT.S3, DTL, ATL, MTAL, CTAL, DCT, PC, IC.A and IC.B. Interestingly, the principal components projection highlights the separation of tubular epithelium from the other cell classes along PC1. Furthermore, PT epithelial cells differentiate from other tubular epithelial cells along PC2. Notably, the positioning of cells along PC2 corresponds to the anatomy of the nephron, with distal cells exhibiting greater separation from PT cells (**Figure 4**).

An important limitation of the single-cell map we constructed, and single-cell technologies in general, is the lack of spatial information. For instance, in scRNAseq analysis, cells are grouped based on transcriptional similarities, disregarding anatomical or spatial context. Thus, to inform the distribution of fructose transporters and metabolic enzymes along the rat nephron, we aimed to establish connections between cell clusters, as defined by the expression of marker genes as well as nephron anatomy. We achieved this in two different ways. Firstly, we assessed the similarity between the average gene expression in scRNAseq clusters and the bulk gene expression in microdissected tubule segments (**Figure 5**). Secondly, we deconvolute bulk transcriptomes from microdissected nephron segments using scRNAseq cellular clusters transcriptomes as a reference. Both approaches yielded consistent evidence for the coexistence of multiple cell types in different subsegment of the nephron. In PTs, both S1 and S3 present more than 75% of their corresponding cell type; however, S2 presents 58% of S2 cells with the remaining cells from adjacent S1 and S3. The abundance of these cells in the S2 segments suggests a differentiation continuum from S1 to S3 with diffused anatomical boundaries. In contrast to PTs, the thin portion of the loop of Henle (DTL and ATL) exhibits high correlation and cell percentages of the respective cell clusters, suggesting high homogeneity of transcriptional phenotypes with well-defined anatomical transitions. We identified two different cell types that map to the thick portion of the loop of Henle, the MTAL phenotype restricted to the MTAL, and the CTAL phenotype present in higher abundance in both subsegments. These cell types may correspond to the medullary “smooth surface” and cortical “rough surface” cells described in rat TAL (38, 39). Microdissected DCT presents a majority of DCT cells expressing NCC (*Slc12a3*); however, DCT transcriptomes not only correlated with the DCT cell cluster but also with those from the TAL. The reason for this is unclear, but it has been reported that the rat TAL could extend beyond the macula densa (40).

Finally, our data indicates that PC is the most abundant cell type in distal tubules, increasing their abundance along CNT, CCD, and OMCD, while IC present an opposite pattern. The PC phenotype was also the most abundant cell type on IMCD. This segment, however, contained 27% of the cells we labeled as “PC.IC”, which were not present in any other segment and may correspond to the previously characterized IMCD cells. Thus, our analysis is consistent with the well-established cell heterogeneity in connecting tubules and the collecting duct system, including the existence of a specialized IMCD cell.

The insights generated from this analysis present limitations inherent to the type and quality of the data. We are also aware that experiments counting a higher number of cells had identified not only more kidney cell types but also different differentiation stages (23). In addition, even though our efforts to identify contaminations with adjacent tubules, it is possible that the microdissected segments contain some cells from other regions. This may explain the correlation between microdissected DCT transcriptomes and TAL cells, as well as the presence of ATL cells in IMCD. Still, with these limitations, we can indicate that the terminology used in scRNAseq cluster assignment referring to specific nephron segments is misleading, as it encompasses anatomical regions with multiple coexisting cell types. This has implications in the way we interpret kidney scRNAseq data as well as for mathematical modelling of nephron transport processes. The consumption of moderately-enriched fructose diets has been associated with the development of salt-sensitive hypertension, in part, due to its actions in the kidney. Thus, once we established the scRNAseq map of the rat kidney, we focused on the study of genes involved in fructose transport and metabolism. We first looked at the polyol pathway, which is the only metabolic pathway known to produce fructose in humans (41). This pathway consists of two enzymes, aldolase reductase (*Akr1b1*), which converts glucose into sorbitol, and sorbitol dehydrogenase (*Sord*) which oxidizes sorbitol to fructose. We found that in both rats and humans, the expression of aldolase reductase is more prominent in DTL, ATL, and IMCD, which is consistent with the necessity to produce sorbitol as an intracellular osmolyte for protecting cells in the hypertonic environment of the inner medulla (**Figure 6**). On the contrary, sorbitol dehydrogenase was primarily expressed in PTs. This suggests that any sorbitol will be rapidly converted to fructose in this segment, thereby preventing sorbitol accumulation and feeding endogenous fructose into metabolic pathways (**Figure 6**). The PT is also responsible for reabsorbing fructose from the luminal fluid. We previously showed that a Na^+^-dependent process drives fructose reabsorption in isolated-perfused rat PTs, as Na^+^ removal from the luminal perfusate reduced fructose reabsorption by 86% (42). Four PT transporters could transport fructose NAGLT1, SGLT4, GLUT5, and SGLT5. Our current analysis showed that the expression of the SGLT5 and GLUT5 genes is significantly higher in the S3 portion of the rat PT as compared to S1 and S2 (**Figure 8**). In addition, SGLT4 was lowly expressed in S2 and S3, while NAGLT1 was highly expressed in S2 and S1. This expression pattern reflects that of human orthologues.

The percentage of ingested fructose that reaches the circulation after the first splanchnic extraction ranges from 15 to 50% (43). Circulating plasma fructose is primarily metabolized by the liver and kidneys (44–47) with the latter accounting for up to 20% of fructose clearance from the plasma (47). Unlike fasting plasma glucose, which is strictly maintained in the mmol/l range by several hormones, fasting fructose concentrations in humans are kept below 20 µmol/l (43). However, ingestion of a fructose-containing meal can raise plasma fructose concentrations more than 15 times, reaching values between ∼30 to 300 µmol/l in humans (43, 48, 49) and ∼50 to 200 µmol/l in mice (45, 50). As such, these concentrations are expected in the early nephron lumen of both humans and rodents. Recent calculations have estimated that, on average, healthy human kidneys filter 4 to 25 g of fructose per day, an amount corresponding to ∼ 8% of the filtered glucose (51). Additionally, fructose produced by the polyol pathway in the kidney medulla could constitute a source of interstitial fructose in the kidney.

NAGLT1, SGLT4, and GLUT5, all have affinities above 1 mmol/l (46, 52–54). The fructose concentration in early PTs likely matches that found in plasma, ranging from 20 to 300 µmol/l (43, 45, 48–50). Thus, even considering the concentrating ability of reabsorbing 70% of fluid, the intratubular fructose concentrations are not expected to rise above 1 mmol/l in any portion of the PT. These expected luminal concentrations of fructose dispute the contribution of these three transporters to the overall fructose reabsorption by the PT epithelia. GLUT5, in particular, has a reported km of 6-9 mmol/l (55), and unlike the other two does not couple Na^+^ to energize transport (55). On the contrary, SGLT5 has an affinity for fructose in the mid-µmol/l range and transports sugars with a 1:1 coupling ratio with Na^+^ (54, 56, 57), fostering the idea that SGLT5 is the largest contributor to the overall reabsorption of fructose. In fact, SGLT5 localizes to the luminal membrane of the rat PT (42), and Slc5a10 (-/-) mice given a fructose-rich diet excrete fructose in the urine (45).

Previous studies (32) show that gene transcripts are, in general, congruent with protein levels. However, protein abundance cannot be predicted from transcript levels only. Therefore, we examined the proteomic expression profile of fructose metabolic enzymes and sugar transporters. We found that the protein expression of both fructose and glucose transporters in microdissected segments were consistent with RNA expression in scRNAseq PT clusters (**Figure 9**). Notably, the expression of SGLT5, fructokinase and triokinase increases from S1 to S3, providing clear evidence that S3 is the main site for fructose reabsorption and metabolism. Together, these data indicate that the transcellular reabsorption of filtered fructose occurs primarily in S3, but with a significant contribution of S2. A more precise estimation, and in particular the contribution of NAGLT1 to this process, would require mathematical modeling including competition with glucose, which is out of the scope of this work. In summary, our data show that under normal conditions, the S3 and S2 segments of the rat PTs are the main site of fructose reabsorption from the tubular lumen.

The scRNAseq data presented here was obtained using a balanced experimental design, which produced data from pools of male and female rats, of different ages subjected to either an ad-libitum or restricted diet regime (20). However, it is not without limitations. An important limitation is the lower number of cells and sequencing depth as compared to newer datasets. A second limitation is that the experimental design does not include metabolic conditions known to alter fructose metabolism. For instance, we previously demonstrated that a diet rich in fructose increases the expression of fructose transporters in rat kidney cortex homogenates (42). This increase correlated with elevated fructose transport rates in the S2 segment, but information in the S3 and S1 segments were missing. In addition, hyperglycemic and hypertonic conditions in diabetes are known to increase aldolase reductase in PTs, which given the constitutive expression of sorbitol dehydrogenase would increase the flux of fructose into metabolic routes. Finally, the SGLT2-inhibitors empagliflozin, dapagliflozin, and remoglipflozin have been shown to also inhibit the human SGLT5 with IC_50_s below 100 µM (54). Even though, the specificity of gliflozins for SGLT2 is higher than for SGLT5, treatment with these drugs may cause a reduction in fructose reabsorption in the pars recta of the PT in at least two ways: 1) by partial inhibition of SGLT5, and 2) by competition with glucose reaching later S2 and S3 segments. Thus, the ability of gliflozins to preserve kidney function (58–60) may be in part due to the reduction of fructose reabsorption in the late PT. Given the significance of fructose metabolism in the development of metabolic syndrome and salt-sensitive hypertension, future studies should investigate fructose reabsorption and metabolism in the kidney, as well as the effects of SGLT2 inhibition during hyperglycemic conditions of consumption of fructose-rich diets.

## Acknowledgments

This work was supported by grants to A.G-V. from the CWRU-SOM Department of Physiology and Biophysics (BGT670107) and the NIH National Institute of Diabetes and Digestive and Kidney Diseases (DK123804).The KPMP is funded by the following grants from the NIDDK: U01DK133081, U01DK133091, U01DK133092, U01DK133093, U01DK133095, U01DK133097, U01DK114866, U01DK114908, U01DK133090, U01DK133113, U01DK133766, U01DK133768, U01DK114907, U01DK114920, U01DK114923, U01DK114933, U24DK114886, UH3DK114926, UH3DK114861, UH3DK114915, UH3DK114937

## Author contributions

Agustin Gonzalez-Vicente conceived the study; Ronghao Zhang, and Agustin Gonzalez-Vicente collected data; Darshan Aatmaram Jadhav, Ronghao Zhang, and Agustin Gonzalez-Vicente analyzed data; Benjamin Kramer, Darshan Aatmaram Jadhav, Ronghao Zhang, and Agustin Gonzalez-Vicente interpreted results; and Ronghao Zhang and Agustin Gonzalez-Vicente prepared the manuscript and figures with input from all authors. All authors provided critical feedback, helping to shape the manuscript. All authors approved the final version of the manuscript.

---

**Supplemental Figure 1:**
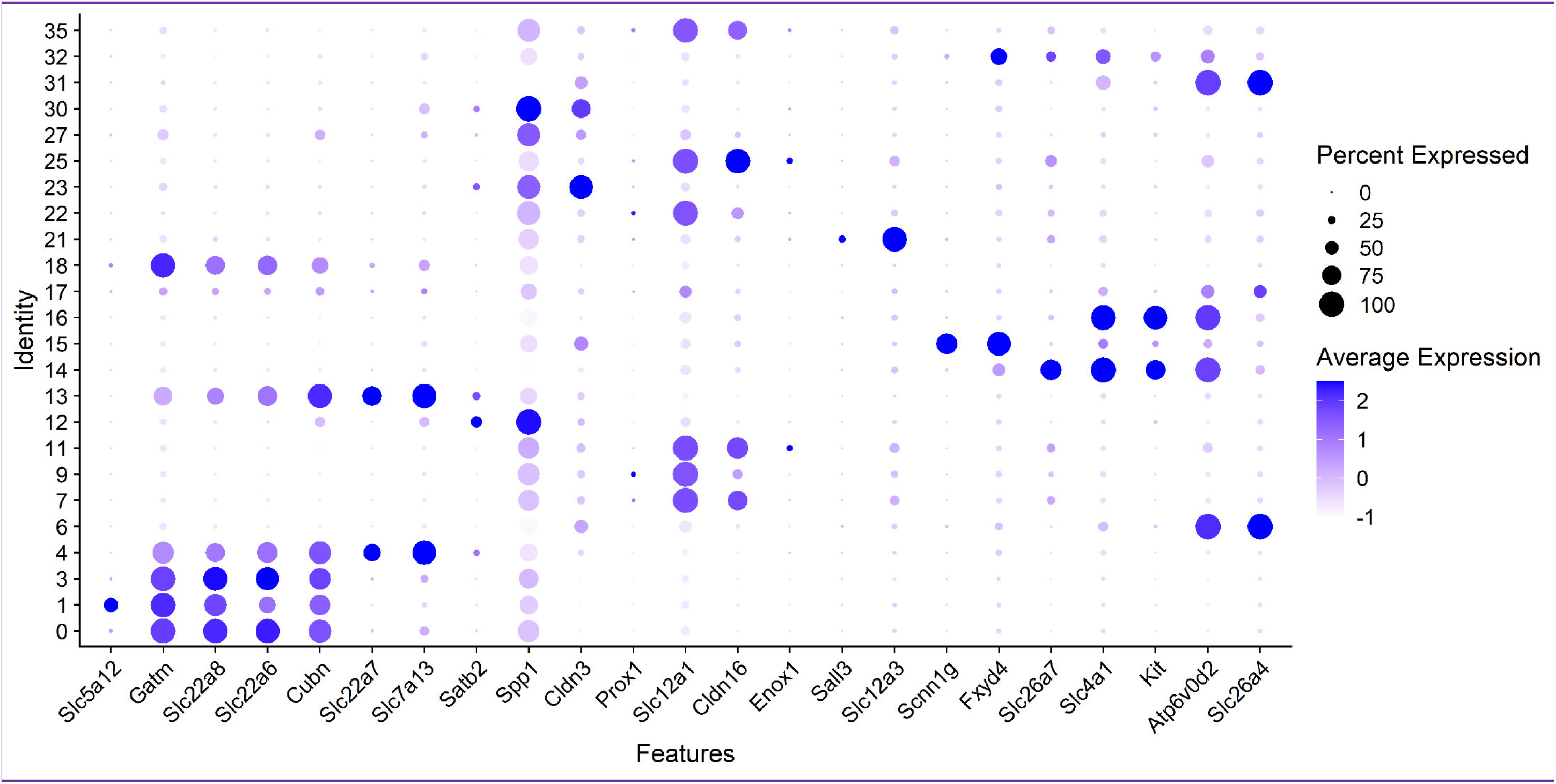
Cluster assignments using HuBMAP Kidney v1.2 cell markers.

**Supplemental Figure 2:**
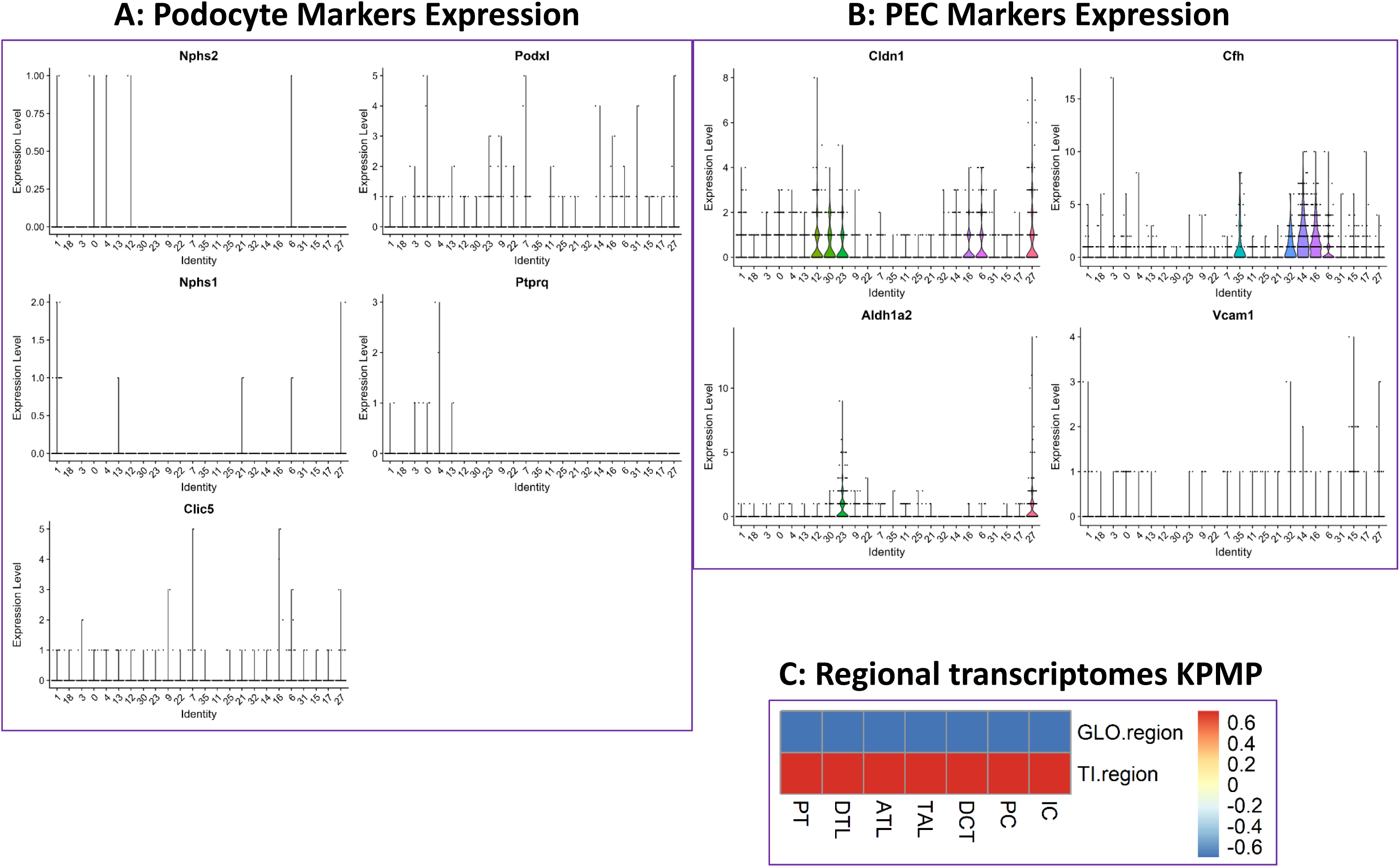
Cluster expression of gene markers for glomerular visceral epithelial cells. **A)** Podocytes and **B)** glomerular parietal epithelial cells (PEC). **C)** Normalized Pearson correlation between rat scRNAseq cell clusters and human regional transcriptomes from the glomerular and tubulointerstitial regions (GLO.region and TI.region, respectively). Publicly available human kidney regional transcriptomics data were obtained from the Kidney Tissue Atlas (atlas.kpmp.org).

**Supplemental Figure 3:**
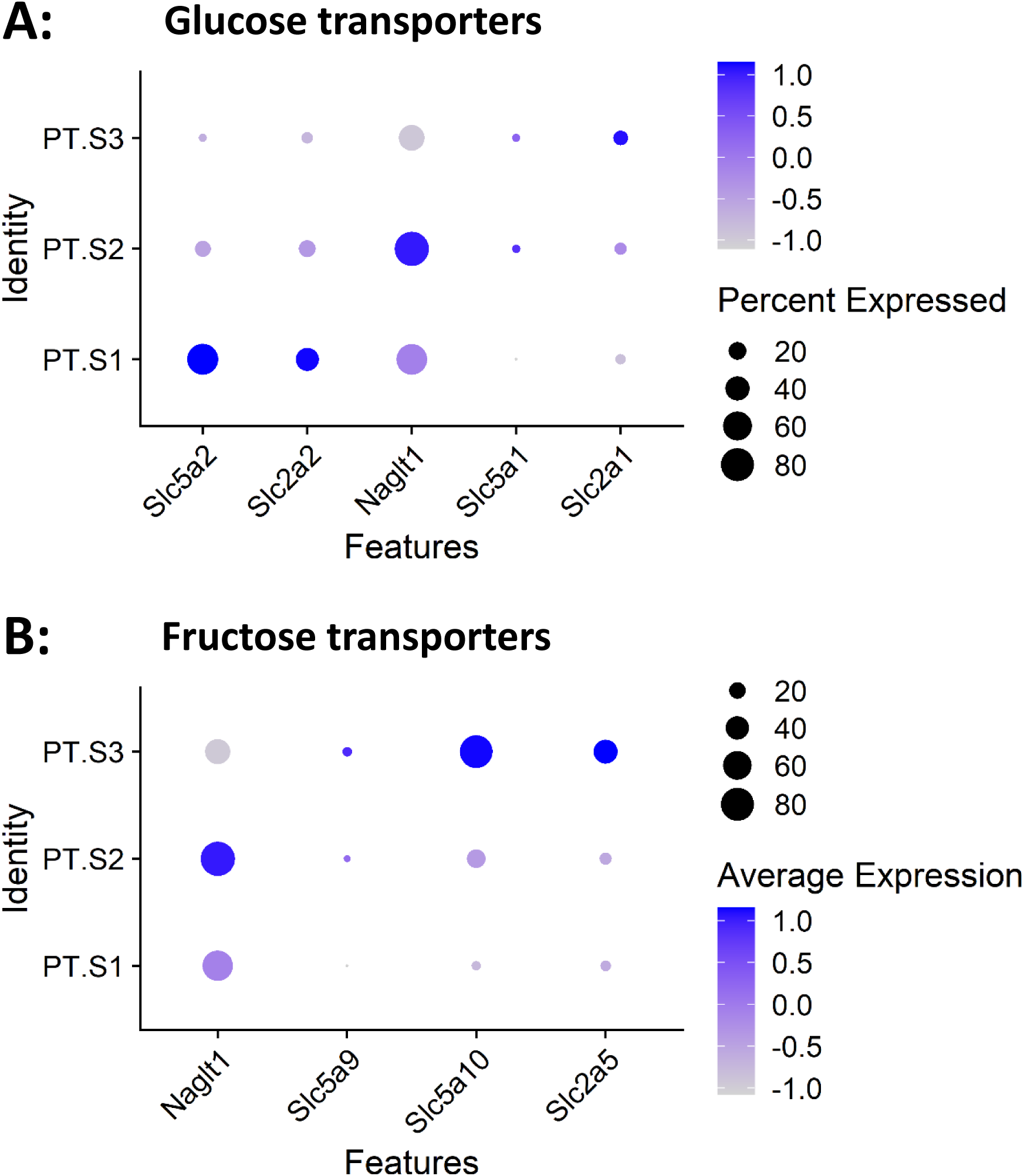
Normalized expression of sugar transporters in rat scRNAseq proximal tubule clusters. **A)** Glucose transporters, and **B)** fructose transporters.

**Supplemental Figure 4:**
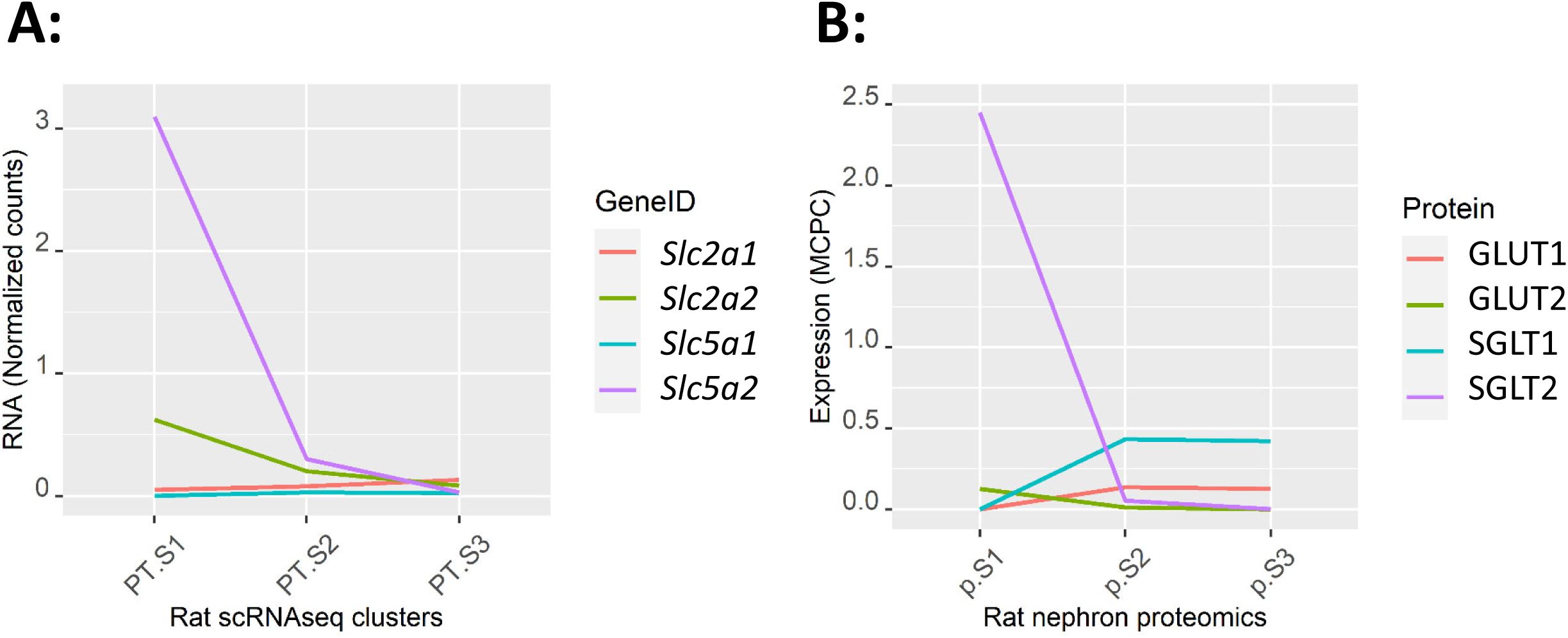
Normalized RNA transcripts of glucose transporters in rat scRNAseq proximal tubule clusters (**A**) compared to their corresponding gene products expressed in “million copies per cell” (MCPC) in microdissected rat proximal tubule segments (**B**). The order and colors of GeneIDs on the left panels match that of the corresponding gene products (Protein) on the right panels. Glucose and fructose transporter NAGLT1 is not shown in this figure, as it has already been shown in Figure 9 of the main manuscript.

**Supplemental Figure 5:**
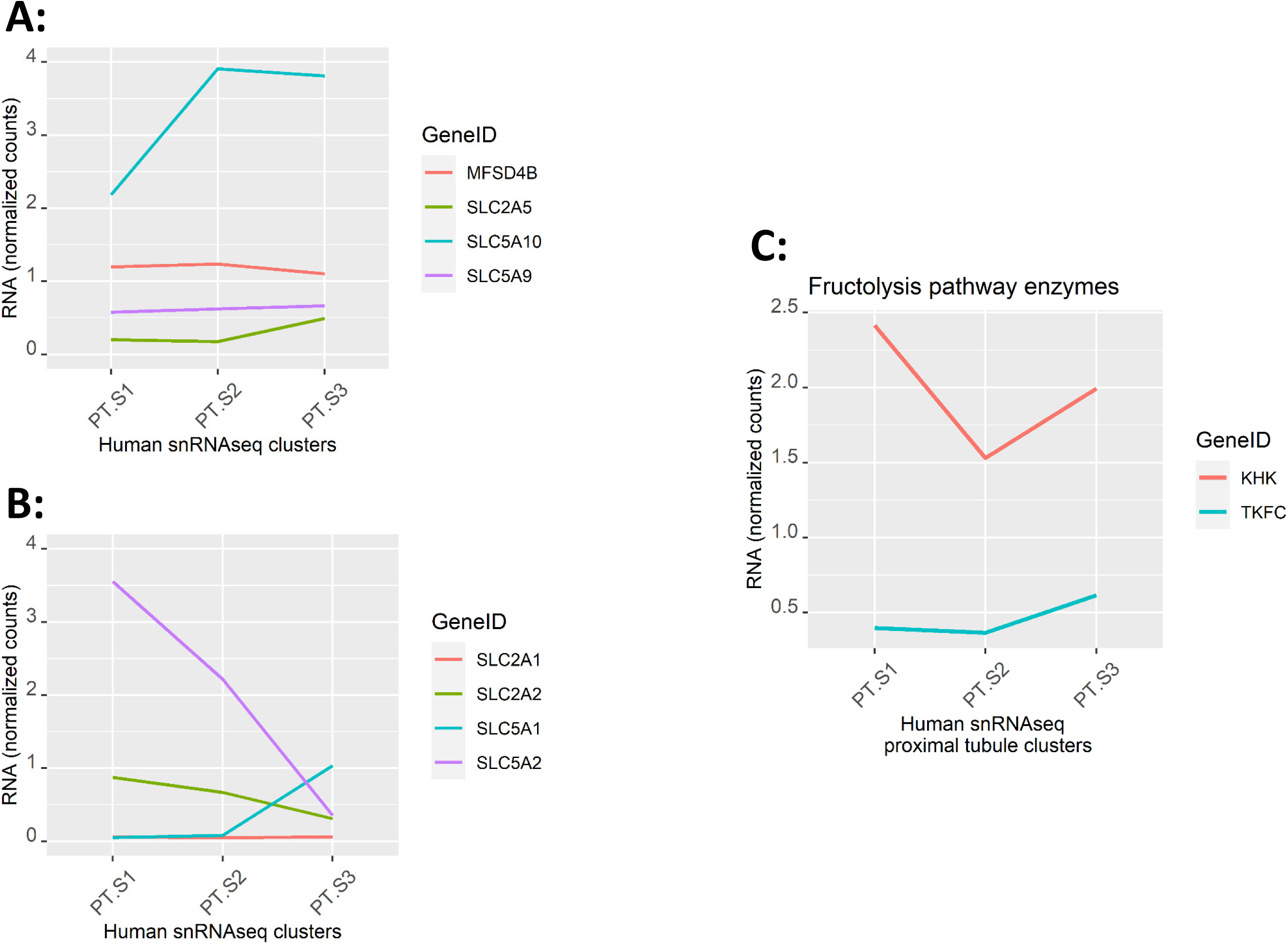
Normalized RNA transcript expression in snRNAseq proximal tubule clusters from human kidneys. **A)** Fructose transporters, **B)** glucose transporters, and **C)** fructokinase (*KHK*) and triokinase (*TKFC*).

**Table S1.**

| Transporter |  | Substrate affinity (km; mmol/l) |  |  | Sugar:Na<sup>+</sup><br>coupling | References |
| --- | --- | --- | --- | --- | --- | --- |
| (Protein) | (Gene) | Glucose | Fructose | Na<sup>+</sup> |  |  |
| SGLT5 | Slc5a10 | 10.1 | 0.62 | 36 | 1:1 | [1, 2, 3] |
| NaGLT1 | Naglt1 | 3.7 | ~6.2 | 202 | 1:1 | [4, 5] |
| GLUT5 | Slc2a5 | - | ~9.3 | - | - | [6] |
| SGLT4 | Slc5a9 | 7.7 | > 10; ~100 (?) | na. | na. | [3, 7] |

**Table S2.** Quality Control.

| Segment.ID | PT | DTL | ATL | TAL | DCT | PC | IC |
| --- | --- | --- | --- | --- | --- | --- | --- |
| S1.1 | 86% | 7% | 0% | 6% | 1% | 0% | 0% |
| S1.2 | 93% | 1% | 0% | 4% | 1% | 0% | 0% |
| S1.3 | 99% | 0% | 0% | 0% | 0% | 1% | 0% |
| S1.4 | 99% | 0% | 0% | 0% | 0% | 0% | 0% |
| S1.5 | 92% | 8% | 0% | 0% | 0% | 0% | 0% |
| S1.6 | 99% | 1% | 0% | 0% | 0% | 0% | 0% |
| S1.7 | 92% | 0% | 0% | 5% | 3% | 0% | 0% |
| S2.1 | 99% | 0% | 0% | 0% | 0% | 0% | 0% |
| S2.2 | 98% | 2% | 0% | 0% | 0% | 0% | 0% |
| S2.3 | 93% | 7% | 0% | 0% | 0% | 0% | 0% |
| S2.4 | 97% | 1% | 0% | 0% | 0% | 2% | 0% |
| S2.5 | 100% | 0% | 0% | 0% | 0% | 0% | 0% |
| S2.6 | 99% | 1% | 0% | 0% | 0% | 0% | 0% |
| S2.7 | 99% | 1% | 0% | 0% | 0% | 0% | 0% |
| S3.1 | 89% | 4% | 0% | 1% | 4% | 2% | 0% |
| S3.2 | 99% | 0% | 0% | 0% | 1% | 0% | 0% |
| S3.3 | 95% | 5% | 0% | 0% | 0% | 0% | 0% |
| S3.4 | 100% | 0% | 0% | 0% | 0% | 0% | 0% |
| S3.5 | 93% | 2% | 0% | 0% | 4% | 0% | 0% |
| S3.6 | 97% | 3% | 0% | 0% | 1% | 0% | 0% |
| S3.7 | 2% | 98% | 0% | 0% | 0% | 0% | 0% |
| SLDTL1 | 0% | 100% | 0% | 0% | 0% | 0% | 0% |
| SLDTL2 | 0% | 57% | 39% | 0% | 0% | 4% | 0% |
| SLDTL3 | 0% | 100% | 0% | 0% | 0% | 0% | 0% |
| SLDTL4 | 0% | 100% | 0% | 0% | 0% | 0% | 0% |
| SLDTL5 | 0% | 100% | 0% | 0% | 0% | 0% | 0% |
| SLDTL6 | 0% | 98% | 0% | 0% | 2% | 0% | 0% |
| SLDTL7 | 0% | 99% | 0% | 0% | 0% | 1% | 0% |
| SLDTL8 | 0% | 99% | 0% | 0% | 0% | 0% | 0% |
| SLDTL9 | 0% | 52% | 48% | 0% | 0% | 0% | 0% |
| SLDTL10 | 0% | 100% | 0% | 0% | 0% | 0% | 0% |
| OMDTL1 | 3% | 94% | 0% | 3% | 0% | 0% | 0% |
| OMDTL2 | 0% | 100% | 0% | 0% | 0% | 0% | 0% |
| OMDTL3 | 0% | 100% | 0% | 0% | 0% | 0% | 0% |
| OMDTL4 | 0% | 100% | 0% | 0% | 0% | 0% | 0% |
| OMDTL5 | 0% | 98% | 0% | 0% | 2% | 0% | 0% |
| OMDTL6 | 0% | 97% | 0% | 1% | 0% | 2% | 0% |
| OMDTL7 | 0% | 100% | 0% | 0% | 0% | 0% | 0% |
| OMDTL8 | 0% | 99% | 0% | 0% | 1% | 0% | 0% |
| OMDTL9 | 0% | 100% | 0% | 0% | 0% | 0% | 0% |
| IMDTL1 | 0% | 99% | 0% | 0% | 1% | 0% | 0% |
| IMDTL2 | 0% | 99% | 0% | 0% | 1% | 0% | 0% |
| IMDTL3 | 0% | 99% | 0% | 0% | 1% | 0% | 0% |
| IMDTL4 | 0% | 100% | 0% | 0% | 0% | 0% | 0% |
| ATL1 | 0% | 0% | 97% | 0% | 0% | 0% | 3% |
| ATL2 | 0% | 0% | 94% | 0% | 0% | 5% | 1% |
| ATL3 | 0% | 0% | 98% | 0% | 1% | 1% | 0% |
| ATL4 | 0% | 99% | 0% | 0% | 1% | 0% | 0% |
| ATL5 | 0% | 7% | 82% | 0% | 11% | 0% | 0% |
| ATL6 | 0% | 100% | 0% | 0% | 0% | 0% | 0% |
| ATL7 | 0% | 99% | 0% | 0% | 1% | 0% | 0% |
| mTAL1 | 0% | 0% | 0% | 99% | 0% | 1% | 0% |
| mTAL2 | 0% | 0% | 0% | 100% | 0% | 0% | 0% |
| mTAL3 | 0% | 0% | 0% | 97% | 0% | 3% | 0% |
| mTAL4 | 0% | 0% | 0% | 99% | 0% | 1% | 0% |
| mTAL5 | 0% | 6% | 0% | 94% | 0% | 0% | 0% |
| mTAL6 | 0% | 35% | 0% | 65% | 0% | 1% | 0% |
| mTAL7 | 0% | 3% | 0% | 97% | 0% | 0% | 0% |
| cTAL1 | 0% | 0% | 0% | 99% | 1% | 0% | 0% |
| cTAL2 | 0% | 0% | 0% | 92% | 8% | 0% | 0% |
| cTAL3 | 0% | 0% | 0% | 95% | 0% | 5% | 0% |
| cTAL4 | 1% | 0% | 0% | 99% | 0% | 0% | 0% |
| cTAL5 | 4% | 0% | 0% | 88% | 7% | 0% | 0% |
| cTAL6 | 0% | 4% | 0% | 95% | 0% | 1% | 0% |
| cTAL7 | 0% | 3% | 0% | 69% | 29% | 0% | 0% |
| DCT.1 | 0% | 2% | 0% | 6% | 92% | 0% | 0% |
| DCT.2 | 0% | 0% | 0% | 0% | 99% | 1% | 0% |
| DCT.3 | 2% | 0% | 0% | 2% | 96% | 0% | 0% |
| DCT.4 | 0% | 0% | 0% | 16% | 80% | 4% | 0% |
| DCT.5 | 0% | 0% | 0% | 2% | 98% | 0% | 0% |
| DCT.6 | 1% | 9% | 0% | 24% | 61% | 6% | 0% |
| DCT.7 | 0% | 10% | 0% | 3% | 87% | 0% | 0% |
| DCT.8 | 0% | 4% | 0% | 3% | 93% | 0% | 0% |
| CNT.1 | 3% | 9% | 0% | 0% | 0% | 35% | 52% |
| CNT.2 | 0% | 7% | 0% | 0% | 10% | 53% | 30% |
| CNT.3 | 0% | 5% | 0% | 0% | 8% | 56% | 31% |
| CNT.4 | 0% | 1% | 0% | 0% | 37% | 14% | 47% |
| CNT.5 | 0% | 3% | 0% | 0% | 13% | 63% | 21% |
| CNT.6 | 0% | 10% | 0% | 0% | 13% | 34% | 43% |
| CCD.1 | 0% | 3% | 0% | 0% | 0% | 89% | 7% |
| CCD.2 | 0% | 3% | 0% | 0% | 0% | 83% | 14% |
| CCD.3 | 0% | 1% | 0% | 0% | 0% | 70% | 29% |
| CCD.4 | 0% | 1% | 0% | 0% | 15% | 64% | 21% |
| CCD.5 | 0% | 6% | 0% | 0% | 6% | 74% | 15% |
| CCD.6 | 0% | 5% | 0% | 0% | 10% | 39% | 46% |
| CCD.7 | 0% | 1% | 0% | 0% | 0% | 98% | 1% |
| CCD.8 | 1% | 2% | 0% | 0% | 2% | 70% | 25% |
| CCD.9 | 0% | 3% | 0% | 0% | 6% | 57% | 34% |
| CCD.10 | 0% | 3% | 0% | 0% | 0% | 71% | 26% |
| CCD.11 | 0% | 4% | 0% | 0% | 11% | 48% | 36% |
| OMCD.1 | 1% | 2% | 0% | 0% | 0% | 72% | 26% |
| OMCD.2 | 0% | 2% | 0% | 0% | 0% | 97% | 1% |
| OMCD.3 | 0% | 1% | 0% | 0% | 0% | 99% | 0% |
| OMCD.4 | 0% | 0% | 0% | 11% | 0% | 72% | 17% |
| OMCD.5 | 0% | 3% | 0% | 0% | 0% | 80% | 17% |
| OMCD.6 | 0% | 1% | 0% | 0% | 1% | 70% | 28% |
| OMCD.7 | 2% | 6% | 0% | 8% | 0% | 70% | 14% |
| IMCD.1 | 0% | 0% | 45% | 0% | 0% | 55% | 0% |
| IMCD.2 | 0% | 0% | 18% | 0% | 0% | 82% | 0% |
| IMCD.3 | 0% | 0% | 22% | 0% | 0% | 78% | 0% |
| IMCD.4 | 0% | 0% | 42% | 0% | 0% | 58% | 0% |
| IMCD.5 | 0% | 0% | 22% | 0% | 0% | 78% | 0% |
| IMCD.6 | 0% | 0% | 38% | 0% | 0% | 62% | 0% |
| IMCD.7 | 0% | 0% | 33% | 0% | 0% | 67% | 0% |
| IMCD.8 | 0% | 0% | 29% | 0% | 0% | 71% | 0% |

DTL from short-loop nephrons are here, but they were not discussed in the main manuscript. The cell proportions are very consistent with those from OMDTL and IMDTL degments.
All IMCD seem to be contaminated with ATL. We did not applied the QC for this segments and mentioned the contamination in the manuscript.

**Table S2a.**
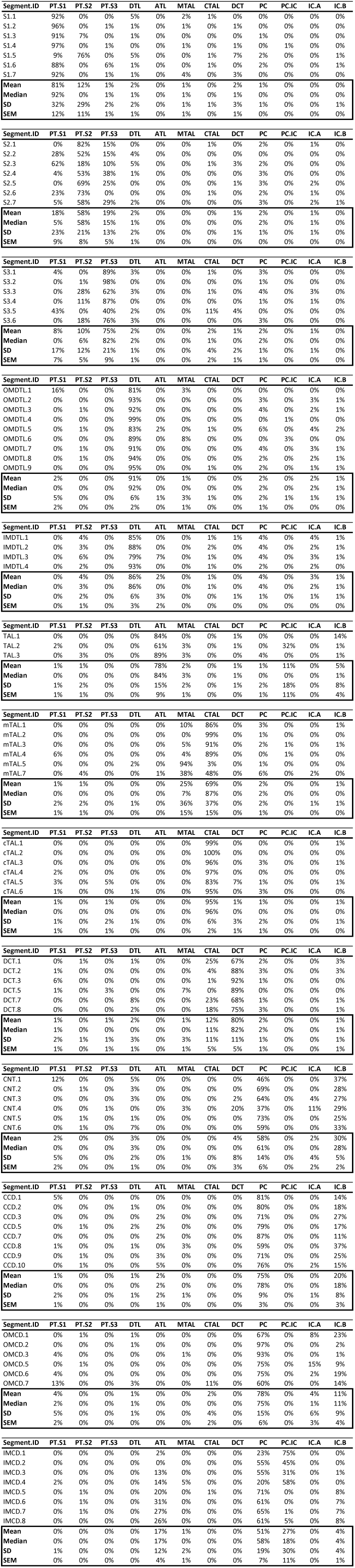
Results.

**Table S2b.** Summary.

| Mean | PT.S1 | PT.S2 | PT.S3 | DTL | ATL | MTAL | CTAL | DCT | PC | IC.A | IC.B | PC.IC |
| --- | --- | --- | --- | --- | --- | --- | --- | --- | --- | --- | --- | --- |
| S1 | 81% | 12% | 1% | 2% | 0% | 1% | 0% | 2% | 1% | 0% | 0% | 0% |
| S2 | 18% | 58% | 19% | 2% | 0% | 0% | 0% | 1% | 2% | 1% | 0% | 0% |
| S3 | 8% | 10% | 75% | 2% | 0% | 0% | 2% | 1% | 2% | 1% | 0% | 0% |
| omDTL | 2% | 0% | 0% | 91% | 0% | 1% | 0% | 0% | 2% | 2% | 1% | 0% |
| imDTL | 0% | 4% | 0% | 86% | 2% | 0% | 1% | 0% | 4% | 3% | 1% | 0% |
| ATL | 0% | 1% | 0% | 0% | 75% | 3% | 0% | 1% | 4% | 2% | 5% | 8% |
| mTAL | 1% | 1% | 0% | 0% | 0% | 25% | 69% | 0% | 2% | 0% | 1% | 0% |
| cTAL | 1% | 0% | 1% | 0% | 0% | 0% | 95% | 1% | 1% | 0% | 1% | 0% |
| DCT | 1% | 0% | 1% | 2% | 0% | 1% | 12% | 80% | 2% | 0% | 1% | 0% |
| CNT | 2% | 0% | 0% | 3% | 0% | 0% | 0% | 4% | 58% | 2% | 30% | 0% |
| cCD | 1% | 0% | 0% | 1% | 2% | 0% | 0% | 0% | 73% | 0% | 23% | 0% |
| omCD | 4% | 0% | 0% | 1% | 0% | 0% | 2% | 0% | 78% | 4% | 11% | 0% |
| imCD | 0% | 0% | 0% | 0% | 17% | 1% | 0% | 0% | 51% | 0% | 4% | 27% |

| Median | PT.S1 | PT.S2 | PT.S3 | DTL | ATL | MTAL | CTAL | DCT | PC | IC.A | IC.B | PC.IC |
| --- | --- | --- | --- | --- | --- | --- | --- | --- | --- | --- | --- | --- |
| S1 | 92% | 0% | 1% | 1% | 0% | 1% | 0% | 1% | 0% | 0% | 0% | 0% |
| S2 | 5% | 58% | 15% | 1% | 0% | 0% | 0% | 0% | 2% | 1% | 0% | 0% |
| S3 | 0% | 6% | 82% | 2% | 0% | 0% | 1% | 0% | 2% | 0% | 0% | 0% |
| omDTL | 0% | 0% | 0% | 92% | 0% | 0% | 0% | 0% | 2% | 2% | 1% | 0% |
| imDTL | 0% | 3% | 0% | 86% | 0% | 0% | 1% | 0% | 4% | 2% | 1% | 0% |
| ATL | 0% | 1% | 0% | 0% | 75% | 3% | 0% | 0% | 2% | 0% | 3% | 0% |
| mTAL | 0% | 0% | 0% | 0% | 0% | 7% | 87% | 0% | 2% | 0% | 0% | 0% |
| cTAL | 0% | 0% | 0% | 0% | 0% | 0% | 96% | 0% | 0% | 0% | 0% | 0% |
| DCT | 1% | 0% | 0% | 0% | 0% | 0% | 11% | 82% | 2% | 0% | 1% | 0% |
| CNT | 0% | 0% | 0% | 3% | 0% | 0% | 0% | 0% | 61% | 0% | 28% | 0% |
| cCD | 0% | 0% | 0% | 1% | 2% | 0% | 0% | 0% | 74% | 0% | 22% | 0% |
| omCD | 2% | 0% | 0% | 1% | 0% | 0% | 0% | 0% | 75% | 1% | 11% | 0% |
| imCD | 0% | 0% | 0% | 0% | 17% | 0% | 0% | 0% | 58% | 0% | 4% | 18% |

**Table S3.**
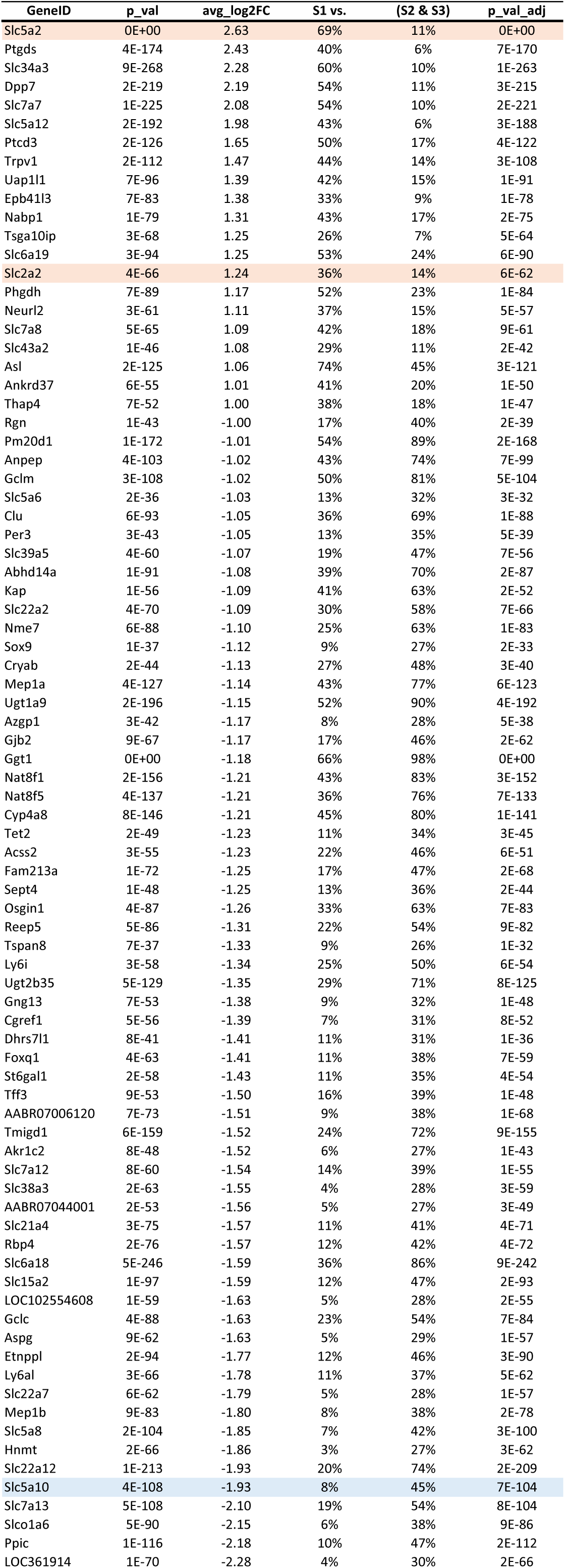
DEG in S1.

**Table S3a.**
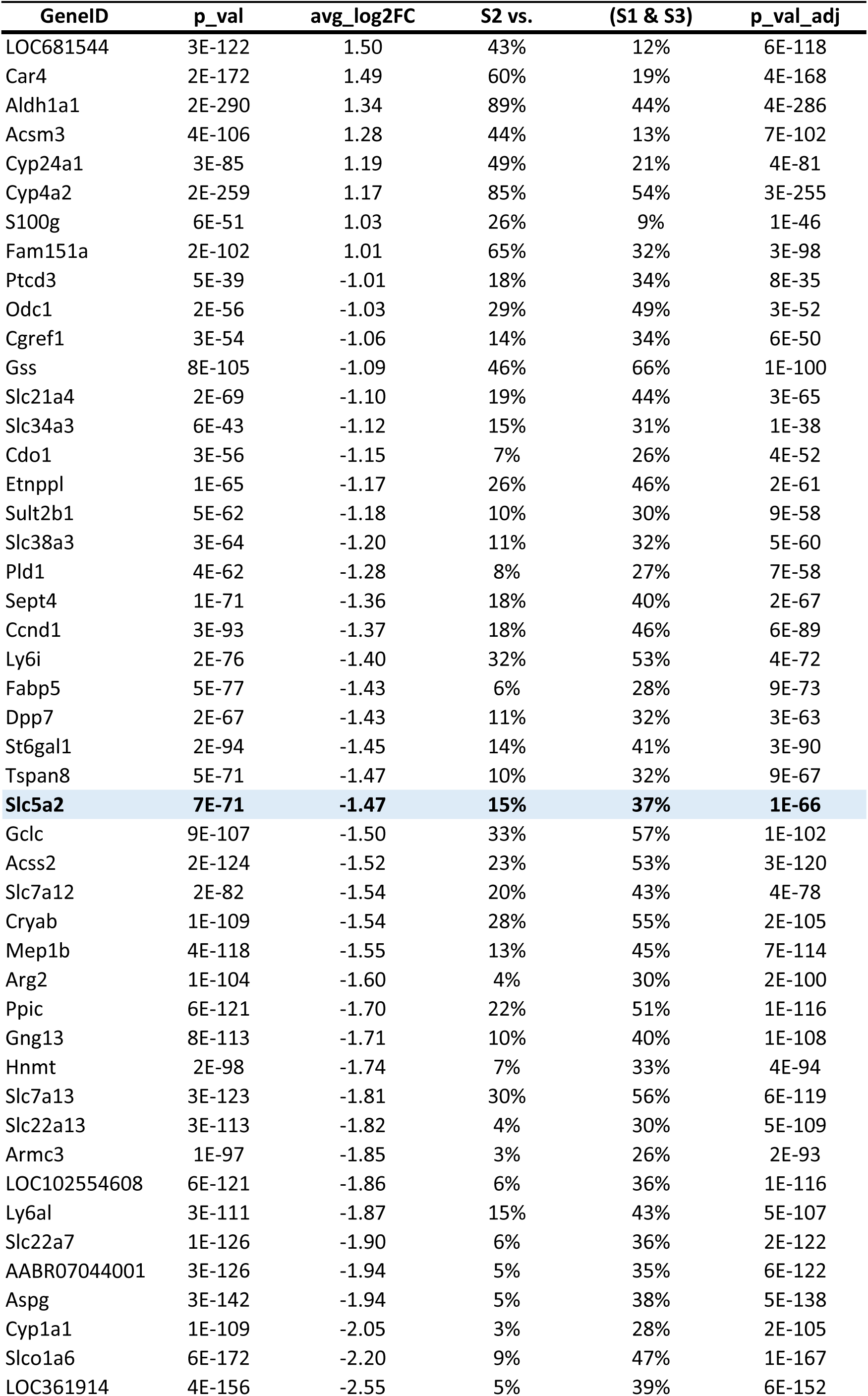
DEG in S2.

| GeneID | p_val | avg_log2FC | S2 vs. | (S1 & S3) | p_val_adj |
| --- | --- | --- | --- | --- | --- |
| LOC681544 | 3E-122 | 1.50 | 43% | 12% | 6E-118 |
| Car4 | 2E-172 | 1.49 | 60% | 19% | 4E-168 |
| Aldh1a1 | 2E-290 | 1.34 | 89% | 44% | 4E-286 |
| Acsm3 | 4E-106 | 1.28 | 44% | 13% | 7E-102 |
| Cyp24a1 | 3E-85 | 1.19 | 49% | 21% | 4E-81 |
| Cyp4a2 | 2E-259 | 1.17 | 85% | 54% | 3E-255 |
| S100g | 6E-51 | 1.03 | 26% | 9% | 1E-46 |
| Fam151a | 2E-102 | 1.01 | 65% | 32% | 3E-98 |
| Ptcd3 | 5E-39 | -1.01 | 18% | 34% | 8E-35 |
| Odc1 | 2E-56 | -1.03 | 29% | 49% | 3E-52 |
| Cgref1 | 3E-54 | -1.06 | 14% | 34% | 6E-50 |
| Gss | 8E-105 | -1.09 | 46% | 66% | 1E-100 |
| Slc21a4 | 2E-69 | -1.10 | 19% | 44% | 3E-65 |
| Slc34a3 | 6E-43 | -1.12 | 15% | 31% | 1E-38 |
| Cdo1 | 3E-56 | -1.15 | 7% | 26% | 4E-52 |
| Etnppl | 1E-65 | -1.17 | 26% | 46% | 2E-61 |
| Sult2b1 | 5E-62 | -1.18 | 10% | 30% | 9E-58 |
| Slc38a3 | 3E-64 | -1.20 | 11% | 32% | 5E-60 |
| Pld1 | 4E-62 | -1.28 | 8% | 27% | 7E-58 |
| Sept4 | 1E-71 | -1.36 | 18% | 40% | 2E-67 |
| Ccnd1 | 3E-93 | -1.37 | 18% | 46% | 6E-89 |
| Ly6i | 2E-76 | -1.40 | 32% | 53% | 4E-72 |
| Fabp5 | 5E-77 | -1.43 | 6% | 28% | 9E-73 |
| Dpp7 | 2E-67 | -1.43 | 11% | 32% | 3E-63 |
| St6gal1 | 2E-94 | -1.45 | 14% | 41% | 3E-90 |
| Tspan8 | 5E-71 | -1.47 | 10% | 32% | 9E-67 |
| **Slc5a2** | **7E-71** | **-1.47** | **15%** | **37%** | **1E-66** |
| Gclc | 9E-107 | -1.50 | 33% | 57% | 1E-102 |
| Acss2 | 2E-124 | -1.52 | 23% | 53% | 3E-120 |
| Slc7a12 | 2E-82 | -1.54 | 20% | 43% | 4E-78 |
| Cryab | 1E-109 | -1.54 | 28% | 55% | 2E-105 |
| Mep1b | 4E-118 | -1.55 | 13% | 45% | 7E-114 |
| Arg2 | 1E-104 | -1.60 | 4% | 30% | 2E-100 |
| Ppic | 6E-121 | -1.70 | 22% | 51% | 1E-116 |
| Gng13 | 8E-113 | -1.71 | 10% | 40% | 1E-108 |
| Hnmt | 2E-98 | -1.74 | 7% | 33% | 4E-94 |
| Slc7a13 | 3E-123 | -1.81 | 30% | 56% | 6E-119 |
| Slc22a13 | 3E-113 | -1.82 | 4% | 30% | 5E-109 |
| Armc3 | 1E-97 | -1.85 | 3% | 26% | 2E-93 |
| LOC102554608 | 6E-121 | -1.86 | 6% | 36% | 1E-116 |
| Ly6al | 3E-111 | -1.87 | 15% | 43% | 5E-107 |
| Slc22a7 | 1E-126 | -1.90 | 6% | 36% | 2E-122 |
| AABR07044001 | 3E-126 | -1.94 | 5% | 35% | 6E-122 |
| Aspg | 3E-142 | -1.94 | 5% | 38% | 5E-138 |
| Cyp1a1 | 1E-109 | -2.05 | 3% | 28% | 2E-105 |
| Slco1a6 | 6E-172 | -2.20 | 9% | 47% | 1E-167 |
| LOC361914 | 4E-156 | -2.55 | 5% | 39% | 6E-152 |

**Table S3b.**
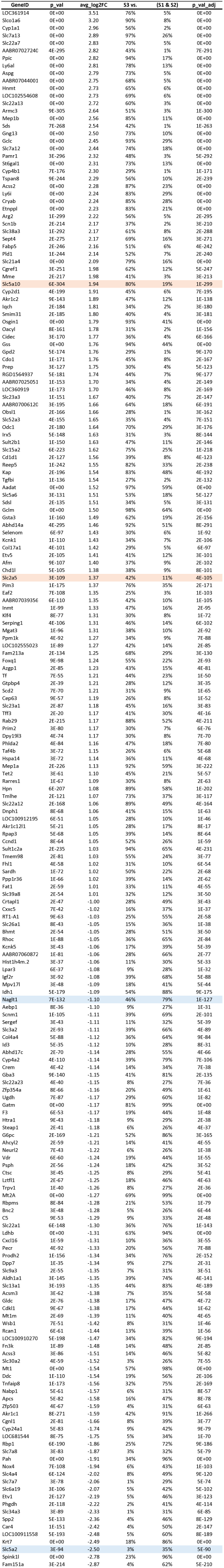
DEG in S3.

## Bibliography

1. Cordain L, Eaton SB, Sebastian A, Mann N, Lindeberg S, Watkins BA, O’Keefe JH, and Brand-Miller J. Origins and evolution of the Western diet: health implications for the 21st century. Am J Clin Nutr 81: 341–354, 2005.

2. National Bureau of Statistics of China http://data.stats.gov.cn/easyquery.htm?cn=C01&zb=A0I0904&sj=2016.

3. China industrial information https://www.chyxx.com/industry/201405/248688.html.

4. Sluik D, Engelen AI, and Feskens EJ. Fructose consumption in the Netherlands: the Dutch National Food Consumption Survey 2007-2010. European journal of clinical nutrition 69: 475–481, 2015.

5. Montonen J, Jarvinen R, Knekt P, Heliovaara M, and Reunanen A. Consumption of sweetened beverages and intakes of fructose and glucose predict type 2 diabetes occurrence. J Nutr 137: 1447–1454, 2007.

6. Vos MB, Kimmons JE, Gillespie C, Welsh J, and Blanck HM. Dietary fructose consumption among US children and adults: the Third National Health and Nutrition Examination Survey. Medscape journal of medicine 10: 160, 2008.

7. Bahadoran Z, Mirmiran P, Tohidi M, and Azizi F. Longitudinal Associations of High-Fructose Diet with Cardiovascular Events and Potential Risk Factors: Tehran Lipid and Glucose Study. Nutrients 9: 872, 2017.

8. Cabral PD, Hong NJ, Hye Khan MA, Ortiz PA, Beierwaltes WH, Imig JD, and Garvin JL. Fructose stimulates Na/H exchange activity and sensitizes the proximal tubule to angiotensin II. Hypertension 63: e68–73, 2014.

9. Gordish KL, Kassem KM, Ortiz PA, and Beierwaltes WH. Moderate (20%) fructose-enriched diet stimulates salt-sensitive hypertension with increased salt retention and decreased renal nitric oxide. Physiological reports 5: 2017.

10. Zenner ZP, Gordish KL, and Beierwaltes WH. Free radical scavenging reverses fructose-induced salt-sensitive hypertension. Integr Blood Press Control 11: 1–9, 2018.

11. Nishimoto Y, Tomida T, Matsui H, Ito T, and Okumura K. Decrease in renal medullary endothelial nitric oxide synthase of fructose-fed, salt-sensitive hypertensive rats. Hypertension 40: 190–194, 2002.

12. Sechi LA. Mechanisms of insulin resistance in rat models of hypertension and their relationships with salt sensitivity. Journal of hypertension 17: 1229–1237, 1999.

13. Catena C, Cavarape A, Novello M, Giacchetti G, and Sechi LA. Insulin receptors and renal sodium handling in hypertensive fructose-fed rats. Kidney Int 64: 2163–2171, 2003.

14. Lewis S, Chen L, Raghuram V, Khundmiri SJ, Chou CL, Yang CR, and Knepper MA. “SLC-omics” of the kidney: solute transporters along the nephron. Am J Physiol Cell Physiol 321: C507–C518, 2021.

15. Diggle CP, Shires M, Leitch D, Brooke D, Carr IM, Markham AF, Hayward BE, Asipu A, and Bonthron DT. Ketohexokinase: expression and localization of the principal fructose-metabolizing enzyme. J Histochem Cytochem 57: 763–774, 2009.

16. Cirillo P, Gersch MS, Mu W, Scherer PM, Kim KM, Gesualdo L, Henderson GN, Johnson RJ, and Sautin YY. Ketohexokinase-dependent metabolism of fructose induces proinflammatory mediators in proximal tubular cells. J Am Soc Nephrol 20: 545–553, 2009.

17. Gonzalez-Vicente A, Garvin JL, and Hopfer U. Transcriptome signature for dietary fructose-specific changes in rat renal cortex: A quantitative approach to physiological relevance. PLoS One 13: e0201293, 2018.

18. Nakagawa T, Johnson RJ, Andres-Hernando A, Roncal-Jimenez C, Sanchez-Lozada LG, Tolan DR, and Lanaspa MA. Fructose Production and Metabolism in the Kidney. J Am Soc Nephrol 31: 898–906, 2020.

19. Lanaspa MA, Ishimoto T, Cicerchi C, Tamura Y, Roncal-Jimenez CA, Chen W, Tanabe K, Andres-Hernando A, Orlicky DJ, Finol E, Inaba S, Li N, Rivard CJ, Kosugi T, Sanchez-Lozada LG, Petrash JM, Sautin YY, Ejaz AA, Kitagawa W, Garcia GE, Bonthron DT, Asipu A, Diggle CP, Rodriguez-Iturbe B, Nakagawa T, and Johnson RJ. Endogenous fructose production and fructokinase activation mediate renal injury in diabetic nephropathy. J Am Soc Nephrol 25: 2526–2538, 2014.

20. Ma S, Sun S, Geng L, Song M, Wang W, Ye Y, Ji Q, Zou Z, Wang S, He X, Li W, Esteban CR, Long X, Guo G, Chan P, Zhou Q, Belmonte JCI, Zhang W, Qu J, and Liu GH. Caloric Restriction Reprograms the Single-Cell Transcriptional Landscape of Rattus Norvegicus Aging. Cell 180: 984–1001 e1022, 2020.

21. Satija_LAB. Cell-Cycle Scoring and Regression https://satijalab.org/seurat/v3.1/cell_cycle_vignette.html.

22. Satija_LAB. Cell cycle genes: 2019 update https://satijalab.org/seurat/reference/cc.genes.updated.2019.

23. Lake BB, Menon R, Winfree S, Hu Q, Ferreira RM, Kalhor K, Barwinska D, Otto EA, Ferkowicz M, Diep D, Plongthongkum N, Knoten A, Urata S, Mariani LH, Naik AS, Eddy S, Zhang B, Wu Y, Salamon D, Williams JC, Wang X, Balderrama KS, Hoover PJ, Murray E, Marshall JL, Noel T, Vijayan A, Hartman A, Chen F, Waikar SS, Rosas SE, Wilson FP, Palevsky PM, Kiryluk K, Sedor JR, Toto RD, Parikh CR, Kim EH, Satija R, Greka A, Macosko EZ, Kharchenko PV, Gaut JP, Hodgin JB, Consortium K, Eadon MT, Dagher PC, El-Achkar TM, Zhang K, Kretzler M, and Jain S. An atlas of healthy and injured cell states and niches in the human kidney. Nature 619: 585–594, 2023.

24. Andrews S. FastQC: A quality control tool for high throughput sequence data. https://github.com/s-andrews/FastQC.

25. Bolger AM, Lohse M, and Usadel B. Trimmomatic: a flexible trimmer for Illumina sequence data. Bioinformatics (Oxford, England) 30: 2114–2120, 2014.

26. Li H, and Durbin R. Fast and accurate short read alignment with Burrows-Wheeler transform. *Bioinformatics (Oxford*, England) 25: 1754–1760, 2009.

27. Danecek P, Bonfield JK, Liddle J, Marshall J, Ohan V, Pollard MO, Whitwham A, Keane T, McCarthy SA, Davies RM, and Li H. Twelve years of SAMtools and BCFtools. Gigascience 10: 2021.

28. Love MI, Huber W, and Anders S. Moderated estimation of fold change and dispersion for RNA-seq data with DESeq2. Genome Biol 15: 550, 2014.

29. Jain; S, Valerius; MT, and Yongqun He. HuBMAP ASCT+B Tables. Kidney v1.2.

30. Kirita Y, Wu H, Uchimura K, Wilson PC, and Humphreys BD. Cell profiling of mouse acute kidney injury reveals conserved cellular responses to injury. Proceedings of the National Academy of Sciences of the United States of America 117: 15874–15883, 2020.

31. Newman AM, Steen CB, Liu CL, Gentles AJ, Chaudhuri AA, Scherer F, Khodadoust MS, Esfahani MS, Luca BA, Steiner D, Diehn M, and Alizadeh AA. Determining cell type abundance and expression from bulk tissues with digital cytometry. Nat Biotechnol 37: 773–782, 2019.

32. Limbutara K, Chou CL, and Knepper MA. Quantitative Proteomics of All 14 Renal Tubule Segments in Rat. J Am Soc Nephrol 31: 1255–1266, 2020.

33. Knepper MA. Epithelial Systems Biology Laboratory https://esbl.nhlbi.nih.gov/Databases/KSBP2/.

34. Lee JW, Chou CL, and Knepper MA. Deep Sequencing in Microdissected Renal Tubules Identifies Nephron Segment-Specific Transcriptomes. J Am Soc Nephrol 26: 2669–2677, 2015.

35. Gonzalez-Vicente A, Pico A, Hanspers K, Slenter D E, and E W. Hexoses metabolism in proximal tubules (WP3916) WikiPathways. https://www.wikipathways.org/pathways/WP3916.html.

36. Gonzalez-Vicente A, Willighagen E, Hanspers K, Slenter D, and E W. Fructose metabolism in proximal tubules (WP3894) WikiPathways. https://www.wikipathways.org/pathways/WP3894.html.

37. Zopf S, Flamig J, Schmid H, Miosge N, Blaschke S, Hahn EG, Muller GA, and Grunewald RW. Localization of the polyol pathway in the human kidney. Histol Histopathol 24: 447–455, 2009.

38. Allen F, and Tisher CC. Morphology of the ascending thick limb of Henle. Kidney Int 9: 8–22, 1976.

39. Mount DB. Thick ascending limb of the loop of Henle. Clin J Am Soc Nephrol 9: 1974–1986, 2014.

40. Yu ASL, Chertow GM, Luyckx VrA, Marsden PA, Skorecki K, and Taal MW. Brenner and Rector’s The Kidney. [Philadelphia, PA]: Elsevier [Philadelphia, PA], 2019.

41. Andres-Hernando A, Johnson RJ, and Lanaspa MA. Endogenous fructose production: what do we know and how relevant is it? Curr Opin Clin Nutr Metab Care 22: 289–294, 2019.

42. Gonzalez-Vicente A, Cabral PD, Hong NJ, Asirwatham J, Saez F, and Garvin JL. Fructose reabsorption by rat proximal tubules: role of Na(+)-linked cotransporters and the effect of dietary fructose. American journal of physiology Renal physiology 316: F473–F480, 2019.

43. Francey C, Cros J, Rosset R, Creze C, Rey V, Stefanoni N, Schneiter P, Tappy L, and Seyssel K. The extra-splanchnic fructose escape after ingestion of a fructose-glucose drink: An exploratory study in healthy humans using a dual fructose isotope method. Clin Nutr ESPEN 29: 125–132, 2019.

44. Laughlin MR. Normal roles for dietary fructose in carbohydrate metabolism. Nutrients 6: 3117–3129, 2014.

45. Fukuzawa T, Fukazawa M, Ueda O, Shimada H, Kito A, Kakefuda M, Kawase Y, Wada NA, Goto C, Fukushima N, Jishage K, Honda K, King GL, and Kawabe Y. SGLT5 reabsorbs fructose in the kidney but its deficiency paradoxically exacerbates hepatic steatosis induced by fructose. PLoS One 8: e56681, 2013.

46. Tazawa S, Yamato T, Fujikura H, Hiratochi M, Itoh F, Tomae M, Takemura Y, Maruyama H, Sugiyama T, Wakamatsu A, Isogai T, and Isaji M. SLC5A9/SGLT4, a new Na+-dependent glucose transporter, is an essential transporter for mannose, 1,5-anhydro-D-glucitol, and fructose. Life Sci 76: 1039–1050, 2005.

47. Bjorkman O, and Felig P. Role of the kidney in the metabolism of fructose in 60-hour fasted humans. Diabetes 31: 516–520, 1982.

48. Pitkanen E. Mannose, mannitol, fructose and 1,5-anhydroglucitol concentrations measured by gas chromatography/mass spectrometry in blood plasma of diabetic patients. Clin Chim Acta 251: 91–103, 1996.

49. Macdonald I, Keyser A, and Pacy D. Some effects, in man, of varying the load of glucose, sucrose, fructose, or sorbitol on various metabolites in blood. Am J Clin Nutr 31: 1305–1311, 1978.

50. Barone S, Fussell SL, Singh AK, Lucas F, Xu J, Kim C, Wu X, Yu Y, Amlal H, Seidler U, Zuo J, and Soleimani M. Slc2a5 (Glut5) is essential for the absorption of fructose in the intestine and generation of fructose-induced hypertension. J Biol Chem 284: 5056–5066, 2009.

51. Vallon V, and Nakagawa T. Renal Tubular Handling of Glucose and Fructose in Health and Disease. Compr Physiol 12: 2995–3044, 2021.

52. Horiba N, Masuda S, Ohnishi C, Takeuchi D, Okuda M, and Inui K. Na(+)-dependent fructose transport via rNaGLT1 in rat kidney. FEBS Lett 546: 276–280, 2003.

53. Uldry M, and Thorens B. The SLC2 family of facilitated hexose and polyol transporters. Pflugers Arch 447: 480–489, 2004.

54. Diederich J, Mounkoro P, Tirado HA, Chevalier N, Van Schaftingen E, and Veiga-da-Cunha M. SGLT5 is the renal transporter for 1,5-anhydroglucitol, a major player in two rare forms of neutropenia. Cell Mol Life Sci 80: 259, 2023.

55. Nomura N, Verdon G, Kang HJ, Shimamura T, Nomura Y, Sonoda Y, Hussien SA, Qureshi AA, Coincon M, Sato Y, Abe H, Nakada-Nakura Y, Hino T, Arakawa T, Kusano-Arai O, Iwanari H, Murata T, Kobayashi T, Hamakubo T, Kasahara M, Iwata S, and Drew D. Structure and mechanism of the mammalian fructose transporter GLUT5. Nature 526: 397–401, 2015.

56. Grempler R, Augustin R, Froehner S, Hildebrandt T, Simon E, Mark M, and Eickelmann P. Functional characterisation of human SGLT-5 as a novel kidney-specific sodium-dependent sugar transporter. FEBS Lett 586: 248–253, 2012.

57. Ghezzi C, Gorraitz E, Hirayama BA, Loo DD, Grempler R, Mayoux E, and Wright EM. Fingerprints of hSGLT5 sugar and cation selectivity. Am J Physiol Cell Physiol 306: C864–870, 2014.

58. Vallon V, and Verma S. Effects of SGLT2 Inhibitors on Kidney and Cardiovascular Function. Annu Rev Physiol 83: 503–528, 2021.

59. Vallon V. Renoprotective Effects of SGLT2 Inhibitors. Heart Fail Clin 18: 539–549, 2022.

60. Verma S, Mudaliar S, and Greasley PJ. Potential Underlying Mechanisms Explaining the Cardiorenal Benefits of Sodium-Glucose Cotransporter 2 Inhibitors. Adv Ther 2023.

